# The human leukemic fusion protein MLL-AF4 promotes autophagy and cell death in the fat body of *Drosophila melanogaster*

**DOI:** 10.64898/2026.07.01.735327

**Authors:** Aina Louise C. Haukeland, Julie Aarmo Johannessen, Nora Rojahn Bråthen, Paul Bergeron, Miriam Formica, Jorrit M. Enserink, Helene Knævelsrud

## Abstract

MLL-rearranged (MLLr) leukemia is an aggressive form of acute leukemia driven by chromosomal translocations fusing MLL with one of more than 100 partner genes, most commonly AF4. We previously showed that expression of the human MLL-AF4 fusion protein in the larval hematopoietic system of *Drosophila melanogaster* promotes hyperproliferation. Here, we report that the same oncogene elicits a strikingly opposite response in the larval fat body, inducing cell shrinkage, autophagy, and caspase-dependent cell death in a manner dependent on the intact fusion protein. Autophagy induction preceded caspase activation, yet the two programs operated in parallel rather than in series. Mechanistically, MLL-AF4-expressing fat body cells displayed elevated AMPK phosphorylation and reduced mTORC1 activity, and depletion of AMPKα abolished caspase activation. Despite producing opposite phenotypes, both tissue-specific responses depended on conserved complex partners, suggesting that MLL-AF4 co-opts a shared mechanism to produce starkly different outcomes depending on cellular context. Uncovering how MLL-AF4 induces apoptosis-like phenotypes in the fat body could potentially be used to rewire leukemic signaling and remove malignant cells.

## Introduction

MLL-rearranged (MLLr) leukemia is an aggressive form of acute leukemia which can arise in both myeloid and lymphoblastic lineages and is associated with poor outcomes (Guarnera et al., 2024). The disease develops due to a reciprocal chromosomal translocation affecting the *MLL* (renamed *KMT2A*) gene on chromosome 11q23. This translocation fuses the N-terminal moiety of MLL with the C-terminus of a fusion partner. In fact, over 100 different fusion partners have been characterized in leukemia, with AF4 (AFF1) being the most common (Meyer et al., 2023). The fusion event gives rise to a fusion protein with the ability to transform hematopoietic cells into leukemic blasts with few or no co-operative mutations (Andersson et al., 2015; Greaves, 2015; Krivtsov & Armstrong, 2007).

Wild-type MLL encodes a histone methyltransferase critical for regulation of key genes during development and hematopoiesis, including a subset of *HOX*-genes, such as *HOXA9*, and *HOX*-cofactors (Milne et al., 2002; Yu et al., 1995). In MLLr leukemia these genes are dysregulated, leading to hematopoietic stem cell expansion and blocked differentiation (Aryal et al., 2023; Faber et al., 2009; Milne et al., 2005; Wang et al., 2011; Yokoyama, 2015). The MLL fusion proteins recruit complex partners such as Menin/MEN1, ENL/MLLT1 and AF9/MLLT3, which contribute to MLLr driven dysregulation of gene expression, consequently promoting transformation (Caslini et al., 2007; Slany, 2020; Takahashi & Yokoyama, 2020). Beyond the essential molecular understanding of the mechanisms induced by MLL fusion proteins, identification of other intracellular pathways that can be targeted in leukemia are of great interest.

One such pathway considered as a possibility to target in a cancer context is autophagy (Klionsky et al., 2021). Macroautophagy (hereafter autophagy) is characterized by formation of double-membraned vesicles termed autophagosomes, which wrap around cytoplasmic components. Autophagosomes fuse with lysosomes to create autolysosomes where the cargo is degraded and the building blocks recycled back to the cytoplasm (Li et al., 2020). In cancer, autophagy has been shown to be two-faced: By ensuring the removal of damaged cellular components and aggregates, autophagy can exert a tumor-suppressive function through preventing cellular damage (Klionsky et al., 2021). On the other side, cancer cells can exploit the protective role of autophagy and use it to shield against various stresses including chemotherapy, as well as aiding cancer cells in meeting energy demands during proliferation and growth (Evangelisti et al., 2015; Klionsky et al., 2021; Li et al., 2020).

Necessary for functional autophagy are autophagy-related (Atg) genes, which have distinct roles throughout the process (Li et al., 2020; Tsukada & Ohsumi, 1993). For instance, the Atg1 (mammalian ULK1) kinase complex regulates autophagy initiation, whereas Atg8 is associated with autophagic membranes until it gets degraded in the autolysosomes, and therefore it is commonly used to detect autophagy (Jipa et al., 2021). Autophagy is further regulated by metabolic sensors such as the mechanistic target of rapamycin complex 1 (mTORC1), which is activated by anabolic signals to suppress autophagy by phosphorylating and inhibiting Atg1 complex components (Saxton & Sabatini, 2017). Another key player in the regulation of autophagy is the energy sensor 5′-adenosine monophosphate-activated protein kinase (AMPK), which upon nutrient deprivation inhibits anabolic processes governed by mTORC1 and promotes autophagy by activating Atg1 (Kim et al., 2011).

We have previously shown that the expression of human MLL-AF4 in the lymph gland, which is the main hematopoietic organ in the *Drosophila* larvae, results in hyperplastic lymph gland growth and an increase in circulating hemocytes (Johannessen et al., 2023). Interestingly, expression of human MLL-AF4 in the fat body of *Drosophila* larvae, which is a functional equivalent of the mammalian liver and adipose tissue (Colombani et al., 2003), yielded a strikingly opposite effect. In this study, we show that MLL-AF4 expression in the fat body leads to growth inhibition and upregulated autophagy coinciding with loss of mTORC1 signaling as well as increased AMPK phosphorylation and caspase activity.

## Results

### Expression of human MLL-AF4 leads to disintegration of the larval fat body

To uncover phenotypes induced by MLL-AF4 in the fat body of *Drosophila melanogaster* 3^rd^ instar larvae, human MLL-AF4 (henceforth MLL-AF4), full-length human MLL, or only the MLL N-terminal or AF4 C-terminal part under UAS-control were expressed using a collagen IV (cg)Gal4 driver, specific for the fat body and hemocytes (Asha et al., 2003; Schmid et al., 2014). The larval fat body is comprised of a monolayer of hexagon-shaped cells that extend throughout the larval body (Xi, 2015). While fat body expressing MLL, the MLL N-terminal or AF4 C-terminal moiety displayed intact cells of similar shape and size, MLL-AF4-expressing fat body cells were not only smaller compared to wild type (wt) and the other transgenes but also exhibited disrupted plasma membrane and loss of the characteristic hexagonal cell shape (Figure 1, A, B) as well as reduced nuclear size (Figure 1, C). The disintegration of the larval fat body was also evident in whole larvae, as the GFP signal was greatly reduced upon MLL-AF4 expression compared to wt and expression of the MLL N-terminus only (Supplementary Figure 1, A, B). Neither full length MLL, the MLL N-terminal nor the AF4 C-terminal portion induced these morphological changes in the fat body, although they were expressed at levels either higher or similar to MLL-AF4 (Supplementary Figure 1, C), indicating that the intact fusion protein is necessary for fat body disintegration.

**Figure 1.**
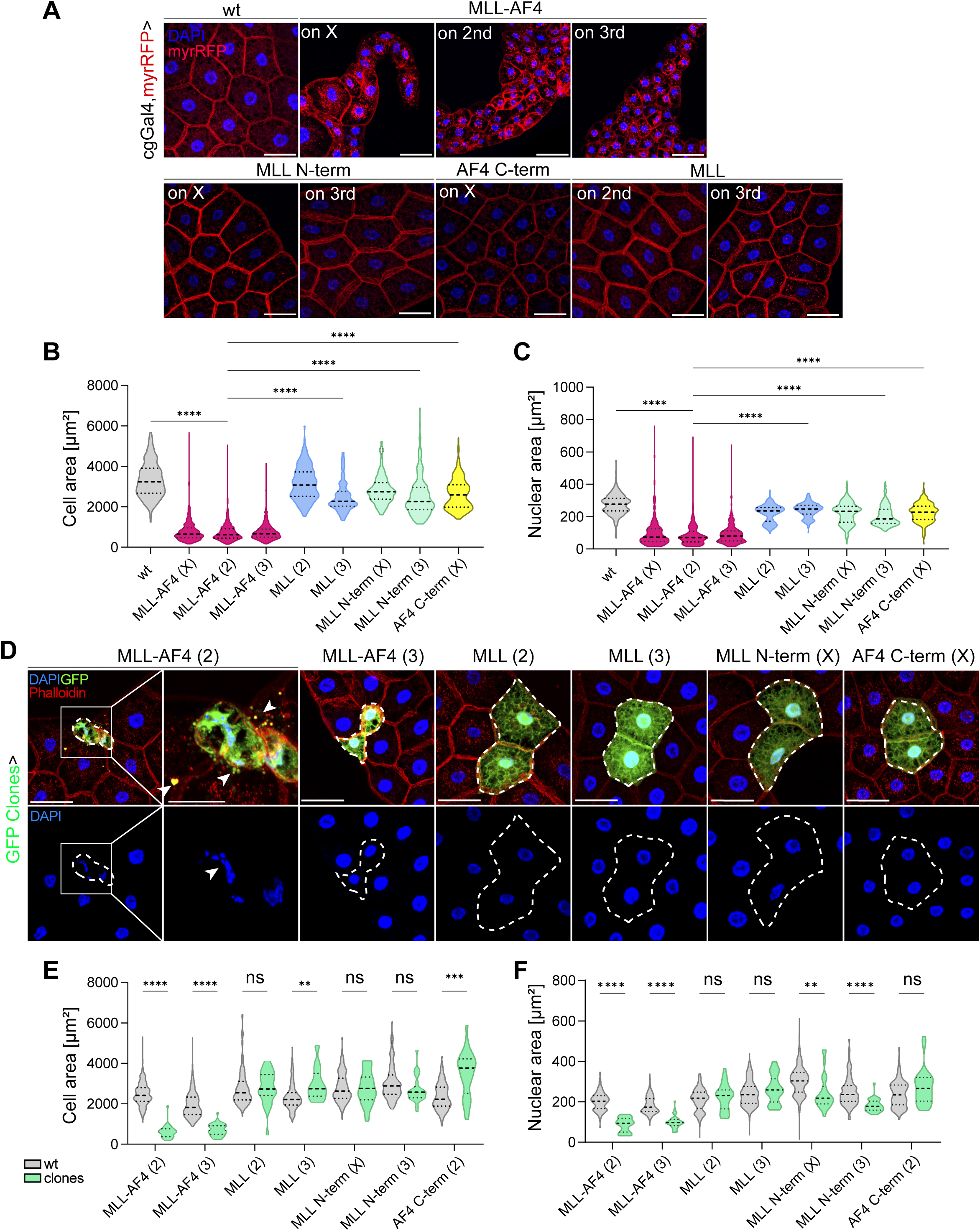
The expression of human MLL-AF4 in the *Drosophila* fat body induces tissue disintegration and cell- autonomously alters fat body morphology. (A) MLL-AF4 drives shrinkage and disintegration of the fat body. Representative confocal fluorescence microscopy images of wL3 larvae that are either wt or expressing respective genotypes and myristoylated RFP driven by cgGal4. Number after genotype refers to the chromosomal location of the transgene. (B-C) Violin plot represents quantification of cell area (B) or nuclear area (C) from images in (A). (D) Cell-autonomous fat body phenotypes by expression of MLL-AF4. Confocal fluorescent images of mosaic fat body of wL3 larvae with Gal4 driven expression of respective transgenes and GFP in clones. Clones are outlined by dotted lines. Fat body is stained with fluorophore conjugated Phalloidin to visualize cellular membranes. Arrowheads in the merged channel annotate membrane blebbing/apoptotic bodies. Arrowhead in the DAPI channel annotates fragmented nuclei. (E-F) Violin plot represents quantification of cell area (E) or nuclear area (F) from images in (D). (B, F) Scale bars = 50 μm. AF4 C-term, AF4 C-terminus. MLL N-term, MLL N-terminus. wt, wild type. wL3, wandering 3rd instar larvae. (B, C, E, F). Genotypes: (A, B, C) wt: *y,w, hsflp/+; cg-Gal4,FRT42D,UAS-myrRFP/+;UAS-GFP-Atg8a/+*, MLL-AF4 (X): *y,w, hsflp/UAS-MLL-AF4; cg-Gal4,FRT42D,UAS-myrRFP/+;UAS-GFP-Atg8a/+*, MLL-AF4 (2): *y,w, hsflp/+; cg-Gal4,FRT42D,UAS-myrRFP/UAS-MLL-AF4;UAS-GFP-Atg8a/+, MLL-AF4* (3): *y,w, hsflp/+; cg-Gal4,FRT42D,UAS-myrRFP/+;UAS-GFP-Atg8a/UAS-MLL-AF4,* MLL (2): *y,w, hsflp/+; cg-Gal4,FRT42D,UAS-myrRFP/UAS- MLL;UAS-GFP-Atg8a/+,* MLL N-term (X): *y,w, hsflp/UAS-MLL N-terminus; cg-Gal4,FRT42D,UAS-myrRFP/+;UAS-GFP-Atg8a/+*, AF4 C-term (X): *y,w, hsflp/UAS-AF4 C-terminus; cg-Gal4,FRT42D,UAS-myrRFP/+;UAS-GFP-Atg8a/+*, (D, E, F) MLL-AF4 (2): *hsflp/+;3xmCherryAtg8a, UAS-GFP/UAS-MLL-AF4;Act>>Gal4, UAS-Dcr2/+.* MLL-AF4 (3): *hsflp/+;3xmCherryAtg8a, UAS-GFP/+;Act>>Gal4, UAS-Dcr2/UAS-MLL-AF4.* MLL (2): *hsflp/+;3xmCherryAtg8a, UAS-GFP/UAS-MLL;Act>>Gal4, UAS-Dcr2/+.* MLL (3): *hsflp/+;3xmCherryAtg8a, UAS-GFP/+;Act>>Gal4, UAS-Dcr2/UAS-MLL.* MLL N-term (X): *hsflp/UAS-MLL N-terminus;3xmCherryAtg8a, UAS-GFP/+;Act>>Gal4, UAS-Dcr2/+.* MLL N-term (3): *hsflp/+;3xmCherryAtg8a, UAS-GFP/+;Act>>Gal4, UAS-Dcr2/UAS-MLL N-terminus.* AF4 C-term (X): *hsflp/UAS-AF4 C-term;3xmCherryAtg8a, UAS-GFP/+;Act>>Gal4, UAS-Dcr2/+*.

### Expression of MLL-AF4 induces cell-autonomous changes to larval fat body cells

To investigate cell-autonomous effects of the oncogene in the fat body, the FLP-FRT system (Golic & Lindquist, 1989) was utilized to express transgenes exclusively in a random subset of cells, indicated as clones. In this mosaic system, the rest of the tissue remains wild-type, enabling direct comparison between cells that express the transgene(s) and wt cells within the same fat body section. Additionally, these clones express UAS-GFP for unambiguous detection. Expression of MLL-AF4 induced cell-autonomous changes in the clones, as these were small, rounded up and showed membrane blebbing or spillage of GFP outside of cell borders (Figure 1, D; indicated by white arrowheads in the merged image) and small or fragmented nuclei (Figure 1, D; indicated by white arrowhead in the DAPI image). These phenotypes are reminiscent of characteristics of caspase-mediated cell death (Mariño et al., 2014; Ramirez & Salvesen, 2018). None of the other transgenes induced these changes, except for a partial reduction of nuclear size in fat body cells expressing the MLL N-terminus (Figure 1, E, F, Supplementary Figure 1, D-F). Moreover, expression of the intact MLL-AF4 oncogene in the fat body increased the number of melanotic masses observed in wL3 larvae (Supplementary Figure 1, G, H).

Together, these data indicate that expression of the intact human leukemic fusion protein MLL-AF4 induces morphological changes reminiscent of cell death in *Drosophila* fat body cells.

### MLL-AF4 promotes caspase activity in the larval fat body cells

Due to several of the MLL-AF4 induced phenotypes coinciding with characteristics of cell death we assessed whether caspases were activated by staining for the active effector caspase cleaved Death caspase-1 (cDcp-1). Indeed, fat body cells expressing the fusion protein exhibited higher cDcp-1 staining intensity compared to the surrounding wt cells, coinciding with the presence of membrane blebbing (yellow arrowhead) (Figure 2, A, B). When expressing MLL-AF4 in the whole tissue, only small, circular cells with disrupted membrane signal (myrRFP) displayed elevated cDcp-1 intensity (yellow arrowheads) (Figure 2, C, D). These results indicate that expression of MLL-AF4 in the fat body promotes caspase-dependent cell death.

**Figure 2.**
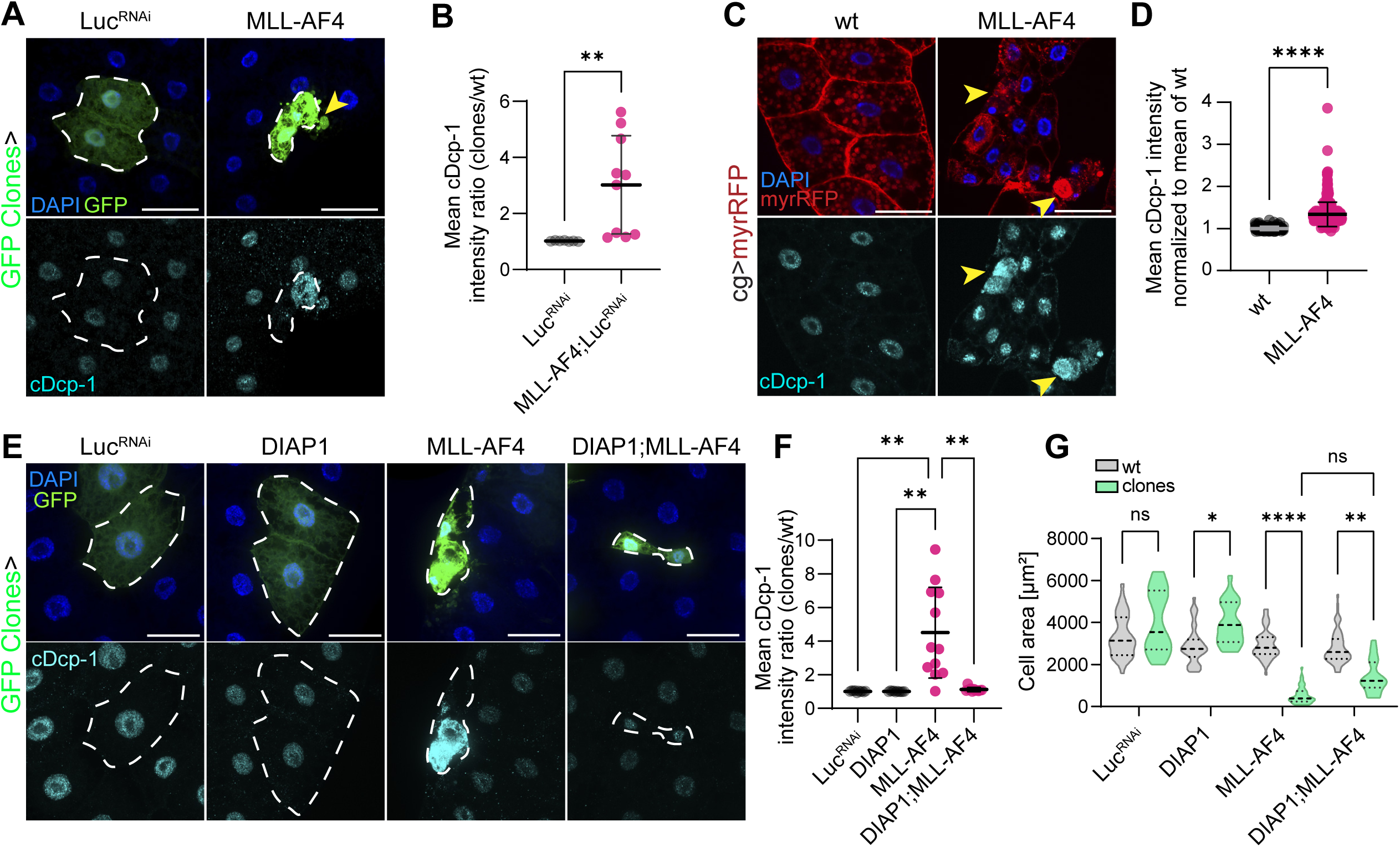
MLL-AF4 promotes caspase activity in fat body cells but caspases are dispensable for MLL-AF4 driven cell size reduction. (A) Representative confocal fluorescence images from mosaic fat body of wL3 larvae expressing either RNAi Luc (control) or MLL-AF4 stained with an antibody against cDcp-1 for caspase activity detection. Yellow arrowhead annotates membrane blob with cDcp-1 staining. (B) Scatter plot with quantification of cDcp-1 intensity in cytosol as a ratio between clones and wt cells. Graph shows mean ± SD. (C) Representative confocal fluorescence images of wL3 fat body that are either wt or expressing human MLL-AF4 with myristoylated RFP expression driven by cgGal4 to visualize membranes. Yellow arrowheads annotate small, round cells with caspase staining. (D) Quantification of mean cDcp-1 intensity in cytosol per cell normalized to wt. Graph shows mean ± SD. (E) Confocal immunofluorescent images of mosaic fat body of wL3 larvae expressing respective transgenes stained with an antibody against cDcp-1 (F) Quantification of mean cDcp-1 intensity ratio in cytosol between clones and wt cells. Graph shows mean ± SD. (G) Violin plot with quantification of cell area of wt and clones for respective genotypes. (A, C, E) Scale bars = 50 μm. cDcp-1, cleaved Death caspase-1. Luc, Luciferase. myrRFP, myristoylated RFP. wt, wild type. Genotypes: (A, B) Luc^RNAi^: *hsflp/+;3xmCherryAtg8a, UAS-GFP/+;Act>>Gal4, UAS-Dcr2/UAS-Luciferase-RNAi TRiP.JF01355.* MLL-AF4: *hsflp/+;3xmCherryAtg8a, UAS-GFP/UAS-MLL-AF4;Act>>Gal4, UAS-Dcr2/+.* (C, D) wt: *y,w, hsflp/UAS-MLL-AF4; cg- Gal4,FRT42D,UAS-myrRFP/+;UAS-GFP-Atg8a/+,* MLL-AF4: *y,w, hsflp/+; cg-Gal4,FRT42D,UAS-myrRFP/UAS-MLL-AF4;UAS-GFP-Atg8a/+.* (E, F, G) Luc^RNAi^: *hsflp/+;3xmCherryAtg8a, UAS-GFP/+;Act>>Gal4, UAS-Dcr2*/*UAS-Dcr2/UAS-Luciferase-RNAi TRiP.JF01355*. DIAP1: *hsflp/+;3xmCherryAtg8a, UAS-GFP/ UAS-DIAP1.H (BL6657); Act>>Gal4, UAS-Dcr2/+.*MLL-AF4: *hsflp/+;3xmCherryAtg8a, UAS-GFP/ VDRCsh60200;Act>>Gal4, UAS-Dcr2/UAS-MLL-AF4.* DIAP1;MLL-AF4: *hsflp/+;3xmCherryAtg8a, UAS-GFP/ UAS-DIAP1.H (BL6657); Act>>Gal4, UAS-Dcr2/ UAS-MLL-AF4*.

Based on the observed caspase activity in MLL-AF4 expressing fat body, we questioned whether the phenotype could be rescued by inhibiting caspase activity. Thus, MLL-AF4 was co-expressed with the anti-apoptotic protein *Drosophila* Inhibitor of Apoptosis 1 (DIAP1) in the mosaic tissue. The cytoplasmic cDcp-1 intensity was reduced in the co-expressing cells (Figure 2, E, F). Similarly, MLL-AF4 expression increased the levels of the initiator caspase DRONC (detected by an antibody targeting mammalian cleaved caspase-3 (Fan & Bergmann, 2010)), which were reduced upon co-expression with DIAP1 (Supplementary Figure 2, A, B). However, the reduction of cell size remained unchanged in presence of ectopic DIAP1 (Figure 2, G and Supplementary Figure 2 C). Co-expressing MLL-AF4 with DIAP1 in the full fat body did lead to a 50% increase in GFP intensity in the whole larvae compared to MLL-AF4 alone, indicating partial restoration of fat body morphology (Supplementary Figure 2, D, E). Again, there was no change in cell area upon DIAP1 co-expression, despite the tissue appearing more intact compared to MLL-AF4 alone (Supplementary Figure 2, F, G). Next, we investigated the effect of co-expressing MLL-AF4 with buffy, an anti-apoptotic Bcl-2 homolog mostly involved in stress-induced cell death (Clavier et al., 2016). Although co-expression of buffy significantly reduced the caspase activity induced by MLL-AF4 (Supplementary Figure 2, H, I), it did not mitigate the oncogene-induced reduction in cell size (Supplementary Figure 2, J), thus mirroring the results obtained with co-expression of DIAP1. Collectively, these results suggest that despite caspases being activated downstream of MLL-AF4, they are not solely responsible for the cell size reduction upon oncogene expression.

### MLL-AF4 upregulates autophagy in larval fat body cells

As both a reduction in cell size and cell death can be observed upon increased catabolism through excessive autophagy (Scott et al., 2007), the autophagy reporter mCherryAtg8a was assessed in the mosaic fat body of wL3 larvae. Indeed, expression of MLL-AF4 elevated the mCherryAtg8a signal relative to the surrounding wt cells and induced an accumulation of bright Atg8a positive structures (Figure 3, A, B). We then assessed whether the increased mCherryAtg8a intensity in MLL-AF4 expressing fat body cells was accompanied by an increase of *Atg8a* transcription. Indeed, MLL-AF4 promoted a significant increase in *Atg8a* transcription relative to the Luc^RNAi^ control (Figure 3, C). To determine whether the striking enhancement of mCherryAtg8a intensity was due to increased induction of autophagy or reduced turnover of autophagic vesicles, we turned to tandem-tagged Atg8a. The pH sensitivity of GFP allows the discrimination of autophagosomes vs autolysosomes since GFP fluorescence is quenched in acidic vesicles, whereas mCherry will fluoresce also in the acidic autolysosome (Kimura et al., 2007). Therefore, high flux will be seen as a high proportion of GFP-negative and mCherry-positive spots, whereas inhibition of autophagosome-lysosome fusion or acidification will be observed as spots positive for both GFP and mCherry. As a control for blocked autophagic flux, the larvae were fed chloroquine (CQ), which prevents autophagosome-lysosome fusion (Mauthe et al., 2018). As expected, fat body from the Luc^RNAi^ control fL3 larvae showed no clear autophagic structures at baseline. However, when the larvae were fed CQ, autophagic vesicles positive for both fluorophores appeared, indicating successful halt of the autophagic flux by the drug (Figure 3, D). To measure autophagic flux, Atg8a spots were quantified as all mCherry-positive spots (total Atg8a spots, Figure 3, E – red bars) and those positive for both GFP and mCherry (Figure 3, E – green bars and Figure 3, D-, observed as yellow spots in the merged channels). MLL-AF4-expressing fat body harboured a striking number of large mCherry-positive structures with imperceptible GFP levels (Figure 3, D, E and Supplementary Figure 3, A). This phenotype was comparable to the increased induction of autophagy observed upon knockdown of Tor utilized as a positive control (Figure 3 D, E), thus indicating that the oncogene promotes autophagy in the *Drosophila* larval fat body.

**Figure 3.**
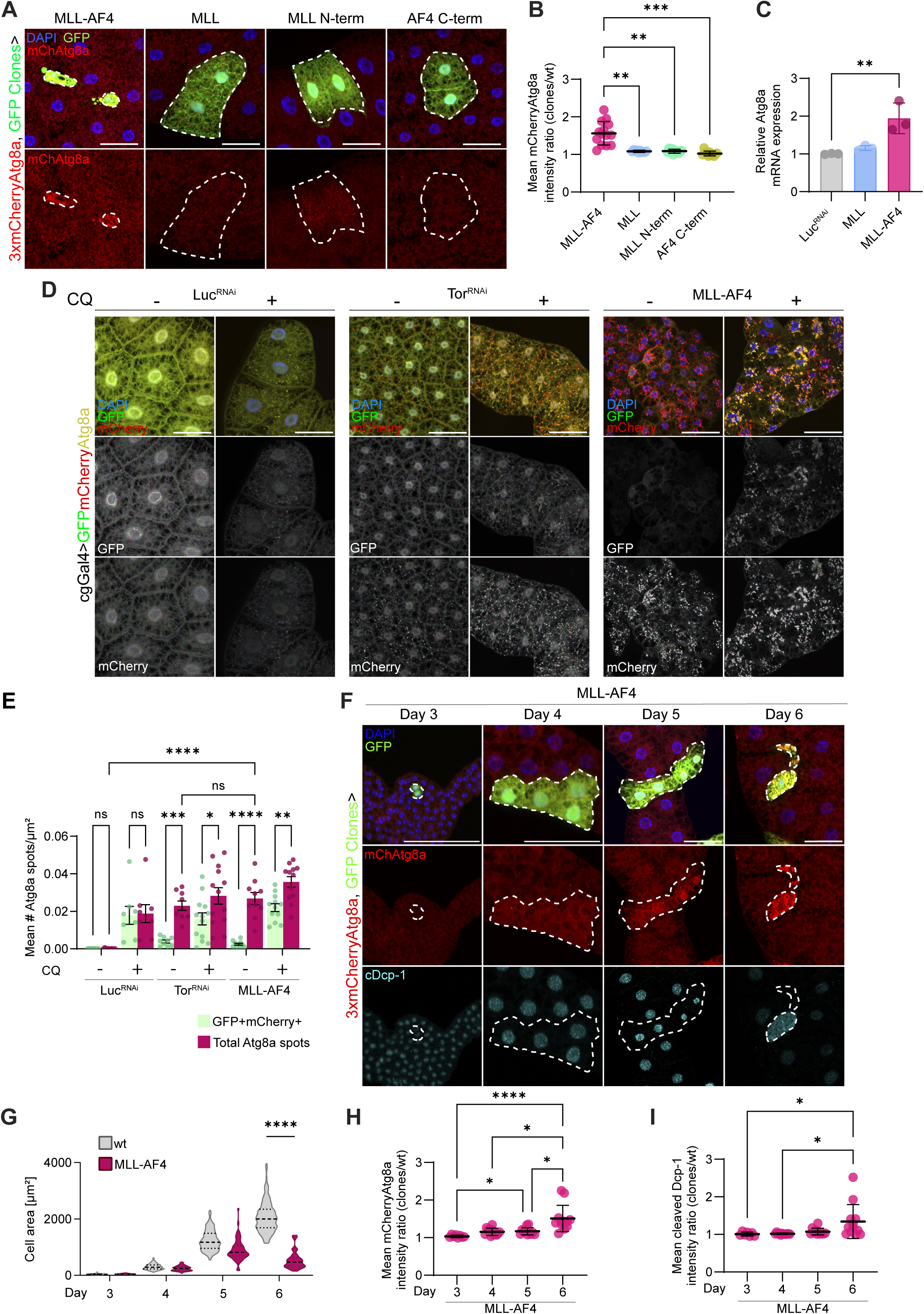
Expression of MLL-AF4 promotes autophagy which precedes caspase activity in fat body cells (A) Representative confocal fluorescent images of mosaic fat body of wL3 larvae expressing respective transgenes in clones. Clones are outlined with a dotted line and express Gal4-driven GFP. The autophagy marker 3xmCherry-tagged Atg8a is expressed in the whole animal under control of the endogenous Atg8a promoter. (B) Quantification of mean mCherry-Atg8a intensity ratio between clones and wt cells. Graph shows mean ± SD. (C) Relative *Atg8a* mRNA expression as determined by RT-qPCR. Graph shows mean ± SD. (D) Representative confocal fluorescent images of fat body of fL3 larvae with cgGal4 driven expression GFP-mCherry-Atg8a co-expressed with indicated transgenes, both non-treated, and fed 15mg/ml chloroquine (CQ) mixed into fly media for 4 hours. (E) Quantification of mean number of both GFP and mCherry positive (green bars) and all Atg8a spots (red bars) per cell area (µm) per image for respective genotypes treated or not with CQ. Bar plot shows mean ± SEM. (F) Representative confocal fluorescent images of mosaic fat body from larvae of different developmental stages indicated by days after egg deposition. Clones expressing human MLL-AF4 are outlined with a dotted line and co-express Gal4-driven GFP. The autophagy marker 3xmCherry-tagged Atg8a is expressed at endogenous levels in the whole animal. The tissue is stained with an antibody against cleaved Death caspase 1 (cDcp-1). (G) Quantification of cell area of wt and clones. Graph shows mean ± SD. (H-I) Quantification of mean intensity ratio of mCherryAtg8a (H) or cytosolic cDcp-1 (I) between clones and wt cells. Graphs show mean ± SD. (A, D, F) Scale bars = 50 µm. (G) Luc, Luciferase. cDcp1-1, cleaved Death caspase-1. CQ, Chloroquine. fL3, feeding 3^rd^ instar. MLL N-term, MLL N-terminus. wL3, wandering 3rd instar. wt, wild type. Genotypes: (A, B) MLL-AF4: *hsflp/+;3xmCherryAtg8a, UAS-GFP/UAS-MLL-AF4;Act>>Gal4, UAS-Dcr2/+*. MLL: *hsflp/+;3xmCherryAtg8a, UAS-GFP/UAS-MLL;Act>>Gal4, UAS-Dcr2/+.* MLL N-term: *hsflp/+;3xmCherryAtg8a, UAS-GFP/+;Act>>Gal4, UAS-Dcr2/UAS-MLL N-terminus.* AF4 C-term: *hsflp/UAS-AF4 C-term;3xmCherryAtg8a, UAS-GFP/+;Act>>Gal4, UAS-Dcr2/+.* (C) Luc^RNAi^: *cgGal4/+; UAS-Luciferase-RNAi TRiP.JF01355/+.* MLL: *cgGal4/UAS-MLL.* MLL-AF4: *cgGal4/UAS-MLL-AF4.* (D, E) Luc^RNAi^: *cg-Gal4, UASp-GFP-mCherry-Atg8a/+;UAS-RNAi Luciferase TRiP.JF01355/+.* mTor^RNAi^: *cg-Gal4, UASp-GFP-mCherry-Atg8a/+;UAS-RNAi mTor TRiP.HMS01114/+.* MLL-AF4: *cg-Gal4, UASp-GFP-mCherry-Atg8a/+;UAS-MLL-AF4/+.* (F, G, H, I) MLL- AF4: *hsflp/+;3xmCherryAtg8a, UAS-GFP/UAS-MLL-AF4;Act>>Gal4, UAS-Dcr2/+*.

### Elevated autophagy driven by MLL-AF4 precedes caspase activity

To understand the relationship between autophagy, caspase activity and cell size, we first dissected mosaic larvae expressing MLL-AF4 in GFP positive clones at different days (3, 4, 5 and 6 after egg deposition) to assess when the decrease in cell size as well as the increase of mCherryAtg8a and caspase activity relative to the surrounding wt cells occurred. There was no significant difference in cell size between MLL-AF4 positive clones and surrounding wt cells until day 6 although there was a tendency to growth inhibition from day 5. Importantly, MLL-AF4 expressing clones tended to have a larger cell size in larvae dissected at day 5 compared to those of day 6, indicating that MLL-AF4 not only restricts cell growth, but also promotes cell shrinkage (Figure 3, F, G). Meanwhile, cells expressing the MLL N-terminal part grew in a similar pattern to that of the surrounding wt cells throughout larval development (Supplementary Figure 3, B, C). MLL-AF4 expressing cells showed significant increase in mCherryAtg8a intensity from day 5, but the most extensive increase occurred on day 6 (Figure 3, H). The cDcp-1 intensity did not significantly increase in MLL-AF4 expressing cells until day 6 (Figure 3, I). Although the expression of the MLL N-terminus displayed slightly elevated mCherryAtg8a intensity throughout development, the cDcp-1 intensity remained equal to that of the surrounding wt cells (Supplementary Figure 3 D, E). Together, these results show that MLL-AF4 expression results in increased autophagic activity in the fat body cells of 3^rd^ instar *Drosophila* larvae and that autophagy precedes caspase activity in this context.

### MLL-AF4 induced cell size reduction and caspase activity is partially dependent on autophagy

As inhibition of caspases failed to rescue the MLL-AF4 induced reduction in cell size, we asked whether inhibition of autophagy could. To investigate this, we depleted MLL-AF4 expressing fat body cells of Atg1. Absence of Atg1 alleviated MLL-AF4 driven cell-autonomous cell shrinkage and led to a significant reduction of mCherryAtg8a intensity (Figure 4, A-C). Depletion of Atg7 did not significantly modify the effect of MLL-AF4 on cell size, but reduced the mCherryAtg8a intensity (Supplementary Figure 4, A-C). However, caspase activity was still detected in absence of these autophagy regulators although there was a tendency of lower cDcp-1 signal compared to MLL-AF4 alone (Figure 4, D, Supplementary Figure 4, D). Co-expression of MLL-AF4 with RNAi against Atg1 in the whole tissue led to complete removal of Atg8a puncta in fL3 larvae (Figure 4, E, F). However, the cell size remained small in absence of autophagy (Figure 4, G). Similar observations were made upon Atg7 depletion (Supplementary Figure 4, E–G). Moreover, caspase activity was detected in Atg1 or Atg7 depleted MLL-AF4 expressing fat body of wL3 larvae although the cDcp-1 signal was significantly lower for Atg7 depleted MLL-AF4 expressing fat body compared to MLL-AF4 alone (Figure 4, H, I, Supplementary Figure 4 H, I). Finally, we investigated whether synchronous inhibition of autophagy and caspase activity could attenuate MLL-AF4 induced phenotypes by co-expressing MLL-AF4 with both DIAP1 and either Atg1 or Atg7 RNAi (Supplementary Figure 4, J). In all of these genotypes, the cell area remained smaller compared to surrounding wt cells (Supplementary Figure 4, K), although clones co-expressing MLL-AF4, DIAP1 and RNAi against Atg1 or Atg7 had a lower mCherryAtg8a spot count (Supplementary Figure 4, L) and the caspase activity was decreased in all conditions with DIAP1 co-expression (Supplementary Figure 4, M). These data indicate that MLL-AF4 driven autophagy contributes to cell size reduction and caspase activity but cannot solely explain these phenotypes.

**Figure 4.**
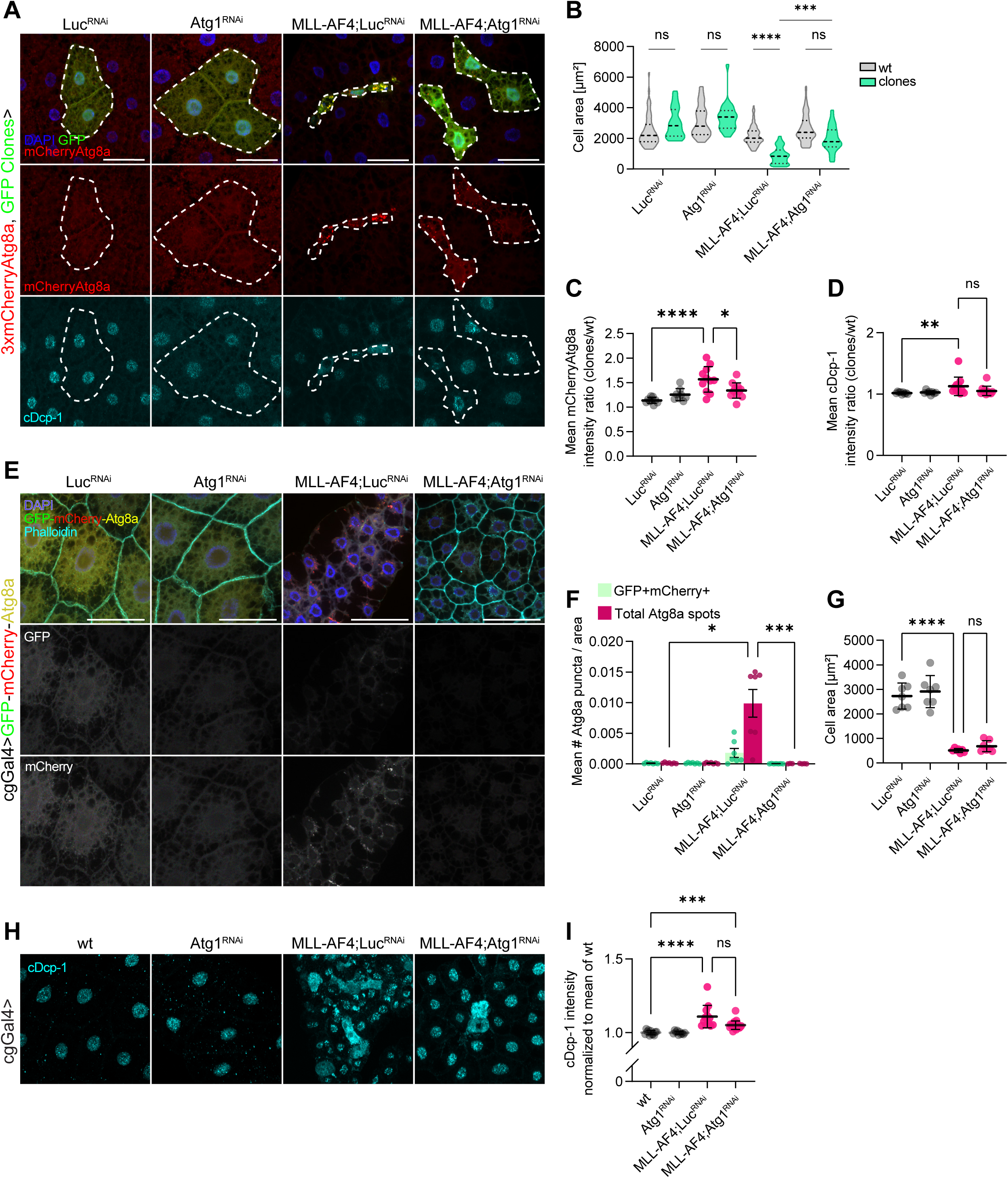
Autophagy is partially responsible for MLL-AF4 driven cell size reduction and caspase activity (A) Mosaic fat body from wL3 with GFP and respective genotypes expressed in clones with endogenous 3xmCherryAtg8a in the whole animal. The tissue is stained with an antibody against cDcp-1 for caspase activity detection. (B) Violin plot shows quantification of cell area from images in (A). (C-D) Quantification of mean mCherryAtg8a (C) or cytosolic cDcp-1 (D). Intensity shown as a ratio between clones and wt cells. (E) Fat body from fL3 larvae expressing cgGal4 driven GFP-mCherry-Atg8a and respective transgenes. The tissue is stained with fluorophore conjugated Phalloidin to visualize cellular membranes. (F) Quantification of Atg8a puncta from images in (E). Red bars represent total Atg8a puncta, while green bars represent Atg8a puncta positive for both GFP and mCherry. Results are presented as mean ± SEM. (G) Quantification of mean cell area per tissue section from images in (E). (H) Fat body from wL3 larvae with cgGal4 driven expression of respective genotypes. The tissue is stained with an antibody against cDcp-1. (G) Quantification of cDcp-1 intensity. (A, E, H) Scale bars = 50 µm. (C, D, G, I) Results are presented as the mean ± SD. A.U., Arbitrary units. cDcp-1, cleaved Death caspase-1. fL3, feeding 3^rd^ instar. Luc, Luciferase. wL3, wandering 3rd instar. wt, wild type. Genotypes: (A, B, C, D) Luc^RNAi^: *hsflp/+;3xmCherryAtg8a, UAS-GFP/+;Act>>Gal4, UAS-Dcr2/UAS-Luciferase-RNAi TRiP.JF01355.* Atg1^RNAi^: *hsflp/+;3xmCherryAtg8a, UAS-GFP/+;Act>>Gal4, UAS-Dcr2/ UAS-Atg1-RNAi TRiP.JF02273.* MLL-AF4;Luc^RNAi^: *hsflp/+;3xmCherryAtg8a, UAS-GFP/UAS-MLL-AF4;Act>>Gal4, UAS-Dcr2/UAS-Luciferase-RNAi TRiP.JF01355.* MLL-AF4;Atg1^RNAi^: *hsflp/+;3xmCherryAtg8a, UAS-GFP/UAS-MLL-AF4;Act>>Gal4, UAS-Dcr2/ UAS-Atg1-RNAi TRiP.JF02273.* (E, F, G) Luc^RNAi^: *cg-Gal4, UASp-GFP-mCherry-Atg8a/+;UAS-RNAi Luciferase TRiP.JF01355/+.* Atg1^RNAi^: *cg-Gal4, UASp-GFP-mCherry-Atg8a/+; UAS-Atg1-RNAi TRiP.JF02273/+.* MLL-AF4;Luc^RNAi^: *cg-Gal4, UASp-GFP-mCherry-Atg8a/UAS-MLL-AF4;UAS-RNAi Luciferase TRiP.JF01355/+.* MLL-AF4;Atg1^RNAi^: *cg-Gal4, UASp-GFP-mCherry-Atg8a/UAS-MLL-AF4; UAS-Atg1-RNAi TRiP.JF02273/+.* (H, I) wt :*y,w, hsflp/+; cg-Gal4,FRT42D,UAS- myrRFP/+;UAS-GFP-Atg8a/+.* Atg1^RNAi^: *y,w, hsflp/+; cg-Gal4,FRT42D,UAS-myrRFP/+;UAS-GFP-Atg8a/ UAS-Atg1-RNAi TRiP.JF02273.* MLL-AF4;Luc^RNAi^: *y,w, hsflp/+; cg-Gal4,FRT42D,UAS-myrRFP/UAS-MLL-AF4;UAS-GFP-Atg8a/ UAS-Luciferase- RNAi TRiP.JF01355.* MLL-AF4;Atg1^RNAi^: *y,w, hsflp/+; cg-Gal4,FRT42D,UAS-myrRFP/UAS-MLL-AF4;UAS-GFP-Atg8a/ UAS-Atg1-RNAi TRiP.JF02273*.

### MLL-AF4 dysregulates Hox-protein Abd-B in the fat body

Hox proteins have been shown to regulate autophagy in *Drosophila* (Banreti et al., 2014), and dysregulation of HOXA9 is a hallmark of MLLr leukemia (Aryal et al., 2023). Therefore, we assessed whether MLL-AF4 led to dysregulation of Hox-proteins in the fat body. Expression levels of the eight *Drosophila Hox*-genes were investigated in MLL-AF4 expressing fat body through qRT-PCR. Strikingly, an approximately twelvefold increase in mRNA levels of *Abd-B*, the *Drosophila* homolog of mammalian *HOXA9* (Argiropoulos & Humphries, 2007), relative to wt was detected in MLL-AF4 expressing fat body (Figure 5, A). Next, Abd-B protein levels were assessed in MLL-AF4 expressing fat body cells, using over-expression of the *Hox*-gene as a control (Figure 5, B). MLL-AF4 increased the presence of Abd-B in both the nuclei and cytoplasm (Figure 5, D, E). Interestingly, over-expression of Abd-B increased mCherryAtg8a intensity (Figure 5, C), although the Hox-protein has been prescribed a repressive role in autophagy regulation (Banreti et al., 2014). Next, we assessed whether depletion of Abd-B could mitigate the MLL-AF4 induced fat body phenotypes (Figure 5, F). Depletion of Abd-B did not increase the cell area, decrease the mCherryAtg8a intensity nor reduce the caspase activity in MLL-AF4 expressing cells (Figure 5, G-I). In fact, the mCherryAtg8a and cDcp-1 intensity was significantly higher in the absence of Abd-B (Figure 5, H, I). Together, these data suggest that MLL-AF4 dysregulates Abd-B expression, but that Abd-B is dispensable for MLL-AF4 induced autophagy and cell death.

**Figure 5.**
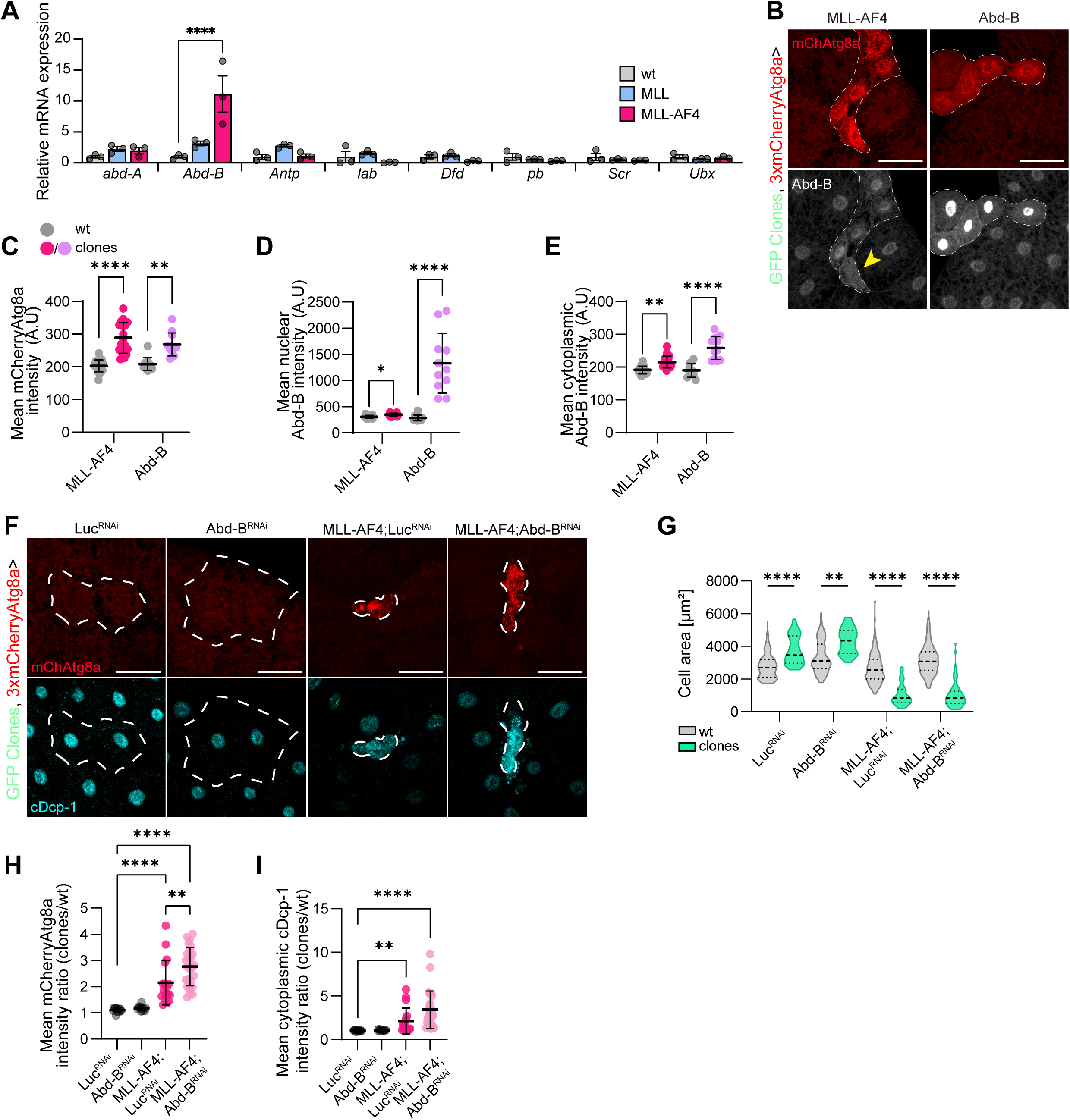
MLL-AF4 upregulates Abd-B expression in the fat body, but Abd-B is not necessary for MLL-AF4 induced fat body phenotypes. (A) Relative mRNA expression of *Drosophila* Hox-genes in the wL3 fat body as determined through qRT-PCR. Results are presented as mean ± SEM. (B) Representative confocal fluorescent images of fat body from wL3 larvae expressing either MLL-AF4 or Abd-B in clones. Clones are outlined with a dotted line. The autophagy marker 3xmCherry-tagged Atg8a is expressed at endogenous levels in the whole animal. The tissue is stained with an antibody against Abd-B. Arrowhead annotates MLL-AF4 expressing cell with no apparent nuclear Abd-B signal but with elevated cytosolic Abd-B. Scale bars = 50 µm. (C) Quantification of mCherryAtg8a intensity of wt cells and clones. (D, E) Quantification of Abd-B intensity in nuclei (D) or cytoplasm (E). (F) Representative confocal fluorescent images of fat body from wL3 larvae expressing respective transgenes in clones. Clones are outlined with a dotted line. The autophagy marker 3xmCherry-tagged Atg8a is expressed at endogenous levels in the whole animal. The tissue is stained with an antibody against cDcp-1. Scale bars = 50 µm. (G) Quantification of cell area of wt and clones shown as a violin plot. (H-I) Quantification of mean mCherryAtg8a (I) or cDcp1 (J) intensity shown as a ratio between clones and wt cells. (C, D, E, H, I) Results are presented as the mean ± SD. A.U., Arbitrary units. cDcp-1, cleaved Drosophila caspase-1. fL3, feeding 3^rd^ instar. wL3, wandering 3rd instar. wt, wild type. Genotypes: (A) wt: *cgGal4/+.* MLL: *cgGal4/UAS-MLL.* MLL-AF4: *cgGal4/UAS-MLL-AF4.* (B-E) MLL-AF4: *hsflp/+;3xmCherryAtg8a, UAS-GFP/UAS-MLL-AF4;Act>>Gal4, UAS-Dcr2/+.* Abd-B: *hsflp/+;3xmCherryAtg8a, UAS-GFP/+;Act>>Gal4, UAS-Dcr2/UAS-Abd-B.* (F-I) Luc^RNAi^: *hsflp/+;3xmCherryAtg8a, UAS-GFP/+;Act>>Gal4, UAS-Dcr2/UAS- Luciferase-RNAi TRiP.JF01355.* Abd-B^RNAi^: *hsflp/+;3xmCherryAtg8a, UAS-GFP/+;Act>>Gal4, UAS-Dcr2/UAS-RNAi Abd-B TRiP.JF02309.* MLL-AF4;Luc^RNAi^: *hsflp/+;3xmCherryAtg8a, UAS-GFP/UAS-MLL-AF4;Act>>Gal4, UAS-Dcr2/UAS-Luciferase-RNAi TRiP.JF01355.* MLL-AF4;Abd-B^RNAi^: *hsflp/+;3xmCherryAtg8a, UAS-GFP/UAS-MLL-AF4;Act>>Gal4, UAS-Dcr2/UAS-RNAi Abd-B TRiP.JF02309*.

### MLL-AF4 complex partners contribute to MLL-AF4 fat body phenotypes

MLLr fusion proteins have been shown to depend on specific complex partners, rendering these complex partners necessary for leukemogenesis (Slany, 2009). We therefore assessed the contribution of MLLr fusion protein complex partners in promoting MLL-AF4 fat body phenotypes. MLL-AF4 was co-expressed with RNAi against Mnn1 or ear, which are *Drosophila* orthologs of MEN1 or ENL/MLLT1 and AF9/MLLT3, respectively. The tissue disintegration was partially or fully rescued when MLL-AF4 was co-expressed with RNAi against Mnn1 or ear (Supplementary Figure 5, A-D). In the cell-autonomous setting, depletion of Mnn1 or ear in MLL-AF4 expressing fat body cells rescued the oncogene driven cell size reduction in fL3 larvae (Supplementary Figure 5E, F). At the wandering stage, depletion of ear mitigated cell shrinkage and Mnn1 followed a similar trend. Both complex partners reduced the mCherryAtg8a intensity (Supplementary Figure 5, G, J) and alleviated the caspase activity (Supplementary Figure 5, K). These results indicate that although the MLL-AF4 fat body phenotype is strikingly opposite of the neoplastic growth observed when the fusion protein is expressed in the hematopoietic tissues of the larvae (Johannessen et al., 2023), it depends on the same conserved complex partners.

Collectively, these results show that MLL-AF4 leads to dysregulation of *Hox*-genes also in the *Drosophila* fat body and that *Drosophila* equivalents of mammalian MLL-AF4 fusion partners are necessary for fat body specific phenotypes induced by the oncogene. However, *Hox*-dysregulation is not necessary for autophagy or cell death promoted by MLL-AF4.

### MLL-AF4 expressing larval fat body cells have low mTORC1 activity coinciding with high AMPK activity

To understand the signalling leading to high autophagy in MLL-AF4 expressing fat body cells, we investigated the activity of mTORC1, by staining the tissue with an antibody against phosphorylated 4E-BP-1 (Battaglioni et al., 2022). In MLL-AF4 expressing cells the high mCherryAtg8a intensity was accompanied with low p4EB-P-1 signal, phenocopying cells depleted of Tor (Figure 6, A-C). Furthermore, the transcription of the *Drosophila* equivalent of *4E-BP-1*, *Thor*, was elevated in MLL-AF4 expressing fat body (Figure 6, D). Whereas active mTORC1 usually represses autophagy, active AMPK promotes autophagy. Accordingly, we observed a significant increase in phosphorylated AMPK in the cytosol of MLL-AF4 expressing fat body cells of fL3 larvae compared to surrounding wt cells. This pAMPK signal was reduced in cells co-expressing MLL-AF4 with RNAi against the catalytic AMPK subunit (AMPKα) (Figure 6, E, F). To further investigate the role of AMPK, we assessed the mCherryAtg8a and cDcp-1 signal in in MLL-AF4 expressing cells depleted of AMPKα at the wL3 stage (Figure 6, G). These cells were smaller than surrounding wt cells, with a tendency to be larger than cells expressing MLL-AF4 alone (Figure 6, H). Furthermore, depletion of AMPK in cells expressing MLL-AF4 resulted in reduced mCherryAtg8a levels and abolished caspase activity (Figure 6, I, J). Collectively, these results indicate that the MLL-AF4 induced autophagy and inhibited cell growth result from increased AMPK and reduced mTORC1 signalling.

**Figure 6.**
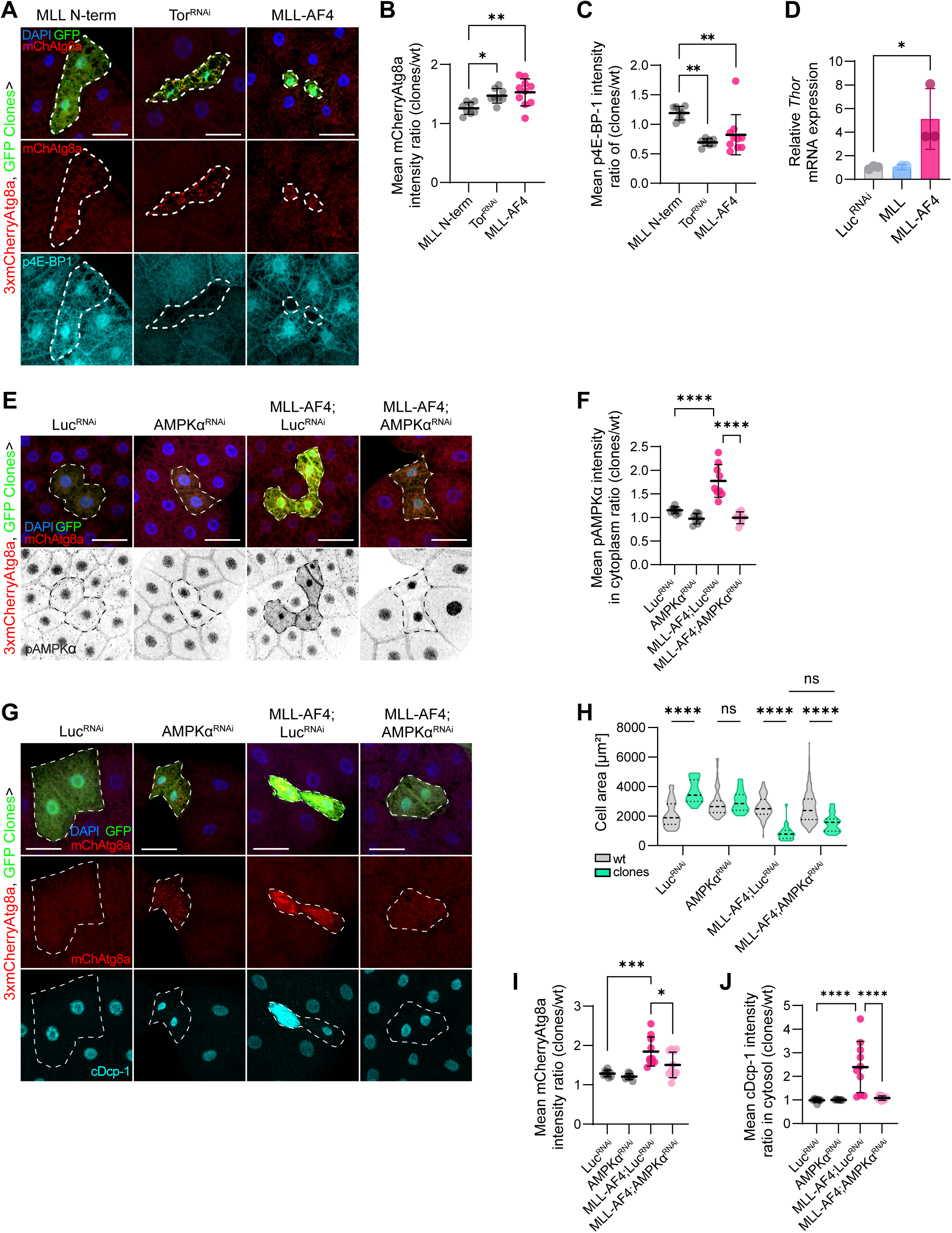
MLL-AF4 expression promotes low mTORC1 activity and high AMPK activity. (A) Representative confocal fluorescent images of fat body from wL3 larvae expressing respective transgenes in clones. Clones are outlined with a dotted line and express Gal4-driven GFP. The autophagy marker 3xmCherry-tagged Atg8a is expressed at endogenous levels in the whole animal. The tissue is stained with an antibody against p4E-BP1. Scale bars = 50 µm. (B-C) Quantification of mean mCherryAtg8a (B) or p4E-BP1 (C) intensity shown as a ratio between clones and wt cells. (D) Relative *Thor* (*Drosophila* 4E-BP1) mRNA expression as determined by RT-qPCR. (E) Representative confocal fluorescent images of mosaic fat body of fL3 larvae expressing respective transgenes in clones. Clones are outlined with a dotted line and express Gal4-driven GFP. The autophagy marker 3xmCherry-tagged Atg8a is expressed at endogenous levels in the whole animal. The tissue is stained with an antibody against pAMPKα. Scale bars = 50 µm. (F) Quantification of mean cytoplasmic pAMPKα intensity shown as the ratio between clones and wt cells. (G) Representative confocal fluorescent images of fat body of wL3 larvae expressing respective transgenes in clones. Clones are outlined with a dotted line and express Gal4-driven GFP. The autophagy marker 3xmCherry-tagged Atg8a is expressed at endogenous levels in the whole animal. The tissue is stained with an antibody against cDcp-1. Scale bars = 50 µm. (H) Quantification of cell area of wt and clones shown as a violin plot. (I-J) Quantification of mean mCherryAtg8a (I) or cDcp-1 (J) intensity shown as a ratio between clones and wt cells. For appropriate panels, results are presented as the mean ± SD. A.U., Arbitrary units. cDcp-1, cleaved Death caspase-1. Luc, Luciferase. fL3, feeding 3^rd^ instar. mChAtg8a, 3xmCherryAtg8a. MLL N-term, MLL N-terminus. wL3, wandering 3rd instar. wt, wild type. Genotypes: (A, B, C) Luc^RNAi^: *hsflp/+;3xmCherryAtg8a, UAS-GFP/+;Act>>Gal4, UAS-Dcr2/UAS-Luciferase-RNAi TRiP.JF01355.* Tor^RNAi^: *hsflp/+;3xmCherryAtg8a, UAS-GFP/+;Act>>Gal4, UAS-Dcr2/ UAS-RNAi Tor TRiP.HMS01114.* MLL-AF4: *hsflp/+;3xmCherryAtg8a, UAS-GFP/UAS-MLL-AF4;Act>>Gal4, UAS-Dcr2/+.* (D) Luc^RNAi^: *cgGal4/+; UAS-Luciferase-RNAi TRiP.JF01355/+.* MLL: *cgGal4/UAS-MLL.* MLL-AF4: *cgGal4/UAS-MLL-AF4.* (E-J) Luc^RNAi^: *hsflp/+;3xmCherryAtg8a, UAS-GFP/+;Act>>Gal4, UAS-Dcr2/UAS-Luciferase-RNAi TRiP.JF01355.* AMPKα^RNAi^: *hsflp/+;3xmCherryAtg8a, UAS-GFP/+;Act>>Gal4, UAS-Dcr2/ UAS-RNAi AMPKalpha TRiP.JF01951.* MLL- AF4;Luc^RNAi^: *hsflp/+;3xmCherryAtg8a, UAS-GFP/UAS-MLL-AF4;Act>>Gal4, UAS-Dcr2/UAS-Luciferase-RNAi TRiP.JF01355.*MLL-AF4;AMPKα^RNAi^: *hsflp/+;3xmCherryAtg8a, UAS-GFP/UAS-MLL-AF4;Act>>Gal4, UAS-Dcr2/ UAS-RNAi AMPKalpha TRiP.JF01951*

## Discussion

Autophagy has contrasting roles in cancer progression. On one hand it is tumor-suppressive, since it ensures removal of damaged cellular components and aggregates, thus preventing a variety of cellular damages. On the other hand, cancer cells can exploit the protective role of autophagy to survive stressful conditions and aid in energy consumption. Moreover, autophagy can help cancer cells meet the elevated energy demands during proliferation and growth (Klionsky et al., 2021; Li et al., 2020). Not surprisingly, studies that investigate the role of autophagy in MLLr leukemia have diverging conclusions. One such study showed that depletion of Atg5 or Atg7 led to more aggressive leukemia progression in mice expressing MLL-ENL (Watson et al., 2015). In concordance, another study showed that Atg5 was essential for efficient initiation of AML in mice models expressing MLL-AF9. However, deletion of Atg5 did not impact the viability of established leukemia stem cells (Q. Liu et al., 2016). Interestingly, Chen et al. reported that MLL-AF9 expressing AML cells experienced high autophagic flux, consistent with the high autophagy flux we observe in MLL-AF4 expressing fat body cells. Furthermore, depletion of Atg5 in the MLL-AF9 expressing AML cells did not affect the growth or survival of these cells (Chen et al., 2017), aligning with how depletion of Atg1 or Atg7 did not inhibit caspase activity in our MLL-AF4 expressing fat body cells. These studies showcase the variety in autophagic responses in MLLr acute leukemia, a reflection of the variability of the disease itself, disease stage and the models used to study it. Normal hematopoiesis occurring in the *Drosophila* larvae has been shown to be dependent on key autophagy players and dysfunctional autophagy leads to aberrant differentiation and proliferation of hemocytes (Katz et al., 2025; Qin et al., 2025). However, little is known of how excessive autophagy affects hematopoiesis in *Drosophila*. The distinct phenotypes observed upon MLL-AF4 expression in the lymph gland or fat body may therefore depend either on inherently different epigenetic profiles between the organs, or on differential effects of Hox-dysregulation and autophagy and should be explored in future studies.

In this study, we observed that expression of MLL-AF4 led to caspase activity and cell death in the wL3 larvae, preceded by elevated autophagy. Overexpression of Atg1 in the fat body upregulates autophagy and induces cell death in a caspase dependent manner (Scott et al., 2007). Conversely, *Drosophila* cell death players such as Hid and Dcp-1 have shown to induce autophagy in the fat body and egg chambers respectively (DeVorkin et al., 2014; Hou et al., 2008; Juhász & Sass, 2005). While it is likely that elevated autophagy drives the cell shrinkage observed upon MLL-AF4 expression in the fat body, inhibition of autophagy did not fully alleviate cell death in this system. Dying cells have been shown to accumulate autophagosomes, as the autophagic flux is impaired to accomplish cell death (Jung et al., 2020). However, accumulation of autophagosomes due to defective flux may also induce cell death (Button et al., 2017). In our system, we have established that the autophagic flux is increased in the MLL-AF4 expressing fat body cells in fL3 larvae. However, we cannot exclude that the elevated levels of mCherryAtg8a observed in wL3 larvae with the highest caspase activity is due to inefficient flux caused by the onset of cell death.

Autophagy and cell death are key mechanisms in the tissue remodeling during *Drosophila* metamorphism. Specifically, during fat body remodeling, autophagy inhibition leads to an increase of caspase activity, and the inhibition of caspases induces upregulation of autophagy (H. Liu et al., 2013). Here, we could not determine any changes in autophagy dependent on caspase inhibition, nor changes in caspase activity upon autophagy inhibition. Therefore, we suggest that MLL-AF4 does not prematurely induce remodeling of the fat body tissue but rather turns on autophagy and cell death programs independently of one another.

Consistent with observations of high autophagic flux, we observed that the MLL-AF4 expressing fat body cells had reduced mTORC1 signaling. Additionally, *Thor* expression was upregulated, which indicates active *Drosophila* forkhead-related transcription factor (FOXO) and results in growth arrest (Jünger et al., 2003; Puig et al., 2003). Together with the high autophagic activity promoted by MLL-AF4, this could possibly explain the reduction in cell size. Moreover, MLL-AF4 expressing fat body cells had elevated AMPK activity, which can inhibit mTORC1 and activate autophagy (Borkowsky et al., 2023; Kim et al., 2011). In murine models of MLL-AF9 leukemia, AMPK has been shown to be crucial for leukemogenesis, potentially by protecting leukemia-initiating cells from metabolic stress. The study further showed that combining depletion of AMPK with dietary restriction had a synergistic effect on AML suppression (Saito et al., 2015). Depleting MLL-AF4 expressing fat body cells of AMPKα did alleviate the autophagic activity as well as caspase activity in wandering 3rd instar larvae. How this is linked to dell death in our system is not completely unraveled. Interestingly, it has been reported that inhibition of AMPK rather promotes cell death in MLLr ALL cells with AMPK hyperactivity (Accordi et al., 2013). Oppositely, it was also found that AMPK can activate p53 mediated cell death in glucose starved mammalian cells (Okoshi et al., 2008) and that activation of AMPK through metformin in ALL cells led to cell growth arrest and UPR-mediated cell death by inducing proteotoxic stress (Leclerc et al., 2013). Notably, AMPK phosphorylation was already elevated in fat body cells during the feeding (fL3) stage, well before caspase activity becomes detectable, demonstrating that elevated AMPK activity is not in itself sufficient to trigger cell death. We propose that the transition to the wandering (wL3) stage acts as a second critical event. Here, abrogation of feeding leads to nutrient deprivation in cells that are already primed by high AMPK activity and elevated autophagy, thus pushing the system past a threshold that engages the caspase machinery. This model is consistent with the temporal data showing autophagy induction at day 5 while caspase activation only becomes significant at day 6 after egg laying, coinciding with the onset of wandering behavior, food withdrawal and developmental autophagy.

Menin is necessary for correct targeting of wild type MLL to chromatin and is crucial for leukemogenesis by MLL fusion proteins (Yokoyama & Cleary, 2008). Hence, Menin inhibitors show promise in clinical trials (Candoni & Coppola, 2024). Consistent with this, the MLL-AF4-induced enlarged lymph gland model was dependent on the *Drosophila* homolog of Menin, Mnn1 (Johannessen et al., 2023). In the fat body, depletion of Mnn1 or ear, the Drosophila ortholog of ENL/MLLT1 and AF9/MLLT3, reduced the tissue disintegration in the whole larvae as well as autophagy and caspase activity in the cell autonomous setting. The dependence on these complex partners indicates that MLL-AF4 might be recruited to and upregulate targets that distinctively promote autophagy and caspase activity in the fat body.

In human MLLr leukemia, ENL/MLLT1 and AF9/MLLT3 are necessary to drive HOXA9 upregulation (Aryal et al., 2023). In our system, MLL-AF4 greatly upregulated mRNA levels of the *Drosophila HOXA9* ortholog *Abd-B*. However, Abd-B protein levels were only slightly increased, and mostly in the cytoplasm, consistent with that Hox-proteins are transported out of the nucleus during late larval stages and degraded through the proteasome (Duffraisse et al., 2020). Interestingly, Abd-B has been characterized as a negative regulator of autophagy in the larval fat body (Banreti et al., 2014). Contradictory to this observation, a recent study showed that co-depletion of Abd-A and Abd-B led to a decrease in expression of autophagy genes in the mid-wandering 3^rd^ instar larvae (Hemba-Waduge et al., 2026). In our system, depletion of Abd-B in MLL-AF4 expressing fat body cells did not mitigate the increased autophagy nor caspase activity but rather aggravated these phenotypes. Furthermore, while forced Abd-B expression in fat body cells did increase the mCherryAtg8a intensity, it did not recapitulate the apoptotic phenotypes induced by MLL-AF4. It is likely that *Hox* dynamics during the developing *Drosophila* larvae complicate the assessment of how Abd-B contributes to MLL-AF4 fat body phenotypes. Taking this into account, we suggest that MLL-AF4 likely dysregulates *Hox*-gene expression in parallel to activating stress signaling in the fat body cells.

Some limitations of the current model should be acknowledged. First of all, we have compared MLL-AF4 to the MLL N-terminal or AF4 C-terminal portions, but we have not assessed the reciprocal fusion protein AF4-MLL, which may be critical for establishing leukemia resulting from t(4;11) translocations (Marschalek, 2020). Secondly, compared to the over-proliferation induced by MLL-AF4 expression in the Drosophila larval hematopoietic system, the cell size reduction and caspase activation observed in the fat body appears as a counterintuitive model to understand human leukemia. However, the fat body phenotype is more robust and easily screenable through the larval cuticle. The large polyploid fat body cells are also better amenable to microscopy-based analysis of intracellular events. We therefore propose to use expression in these two organ systems as complementary models to investigate genetic modifiers of MLL-AF4 phenotypes.

Overall, this study validates that the *Drosophila* larval fat body can be utilized to investigate pathways downstream of MLL-AF4 expression. Further identification of how this leukemia-inducing oncogene promotes apoptosis-like phenotypes in the fat body can be utilized to rewire pro-tumorigenic leukemia signaling into anti-tumor defense signaling.

## Materials and methods

### Drosophila strains

The following Drosophila melanogaster strains were used: wiso14 as a control (Gift from Dr. M. Therrien, University of Montréal). w; UAS-MLL-AF4 ML5 (on 2nd), w; UAS-MLL-AF4 ML6 (on 3rd), UAS-MLL-AF4 ML7 (on X), w; UAS-MLL FL (on 2nd), w; UAS-MLL FL (on 3rd), w; UAS-MLL N-term (on 3rd), w; UAS-MLL N-term (on X), w; UAS-AF4 C-term (on X) (gifts from Dr. R. Paro). The following driver lines were used for clonal or tissue specific expressions: y,w, hs-flp; cg-Gal4,FRT42D,UAS-myrRFP;UAS-GFP-Atg8a (gift from Dr. T. Neufeld), hsflp; 3xmCherry-Atg8a, UAS-GFP; Act>CD2>Gal4, UAS-Dicer2 (gift from Dr. G. Juhasz).

The following strains were acquired from the Bloomington Drosophila stock center (BDSC): w[1118]; P{w[+mC]=Cg-GAL4.A}2 (BDSC #7011), y[1] w[1118]; P{w[+mC]=UASp-GFP-mCherry-Atg8a}2 (BDSC #37749), y[1] v[1]; UAS-Luciferase-RNAi TRiP.JF01355 (BDSC #31603) (used as an RNAi control), w[1]; P{w[+mC]=UAS-Abd-B.m.C}1.1 (BDSC #913), UAS-RNAi AbdB TRiP.JF02309}attP2, UAS-RNAi AMPKalpha TRiP.JF01951}attP2 (BDSC #25931), UAS-Atg1-RNAi TRiP.JF02273 (BDSC #26731), UAS-Atg7-RNAi TRiP.JF02787 (BDSC #27707), w[*]; P{w[+mC]=UAS-Buffy.Q}2 (BDSC #58358), w[*]; P{w[+mC]=UAS-DIAP1.H}3 (BDSC #6657), UAS-RNAi ear TRiP.HMS00107}attP2 (BDSC #34798),, UAS-RNAi Mnn1 TRiP.GL00018}attP2 (BDSC #35150), UAS-RNAi Tor TRiP.HMS01114}attP2 (BDSC #34639)

The following strains were obtained from the Vienna Drosophila RNAi Center (VDRC): 40D-UAS #60101, w[1118];P{VDRCsh60200}attP40 #60200 (both as controls for co-expression with MLL-AF4 on 3rd chromosome).

The fly food used for all crosses was comprised of standard potato mash fly food (27.3 g/liter dry yeast, 32.7 g/liter dried potato powder, 60 g/liter sucrose, 0.73% agar, 0.2% 4-hydroxybenzoic acid methyl ester (nipagin) in 100% ethanol and 0.45% propionic acid). The crosses were incubated at 25°C until dissection/imaging.

### Fluorescence imaging of whole larvae

To visualize phenotypic changes in the larval fat body, fluorescent microscopy images of entire larvae were captured using a GFP filter on either the fluorescence stereoscope Leica Thunder equipped with 1x objective, monochrome camera with a sCMOS sensor, LED light source and filters: 500-550 nm, using the Leica Application Suite software or Leica MZFLIII stereomicroscope and LAS v4.9 software. Larvae were rinsed in 1XPBS before they were dried on fine paper tissue and moved to 70% glycerol on a microscope slide. Larvae were then heat fixed on a 70°C heating block for 10 seconds. Using the microscope, the larvae were arranged and imaged. Post imaging enhancement of the GFP signal for satisfactory visualization and quantification of the GFP signal was performed in ImageJ.

### Chloroquine treatment of *Drosophila* larvae

Chloroquine (CQ) diphosphate salt (Sigma-Aldrich, C6628) was dissolved in distilled water to a concentration of 150 mg/ml. Next, 100 µl of dissolved CQ was mixed with 900 µl of fly food yielding a final concentration of 15 mg/ml. Feeding third instar larvae were moved from vials containing standard fly food and washed in 1XPBS before they were moved into CQ-containing fly food. They were exposed to CQ for 4 hours before dissection.

### Counting melanotic masses

Wandering L3 larvae of each genotype of interest were taken from their tubes with tweezers and then washed twice in PBS with 0,2 % Triton X-100 (0.2%PBT) and once in 1XPBS. The LEICA MZ FL III microscope set at 5x zoom was used to observe the larvae in 1XPBS. Melanotic masses in each individual larva were manually counted while rotating the larva to get a full overview.

### Fluorescence confocal microscopy of fat body from dissected larvae

Fluorescent confocal microscopy was employed to visualize tissue with transgenic expression of fluorescent proteins, or tissues stained with antibodies coupled to fluorophores, or a combination of both. Before dissection, larvae were rinsed twice in 0.2% PBT, then rinsed once in PBS. Between 6-15 larvae per genotype were dissected by pinching a hole in the cuticle and inverting the animal to expose the fat body. The carcasses were then fixed in 200 μl of 4% paraformaldehyde (Polysciences #18814-20) in 1XPBS at a temperature of 4 °C overnight or at room temperature for 1 hour. Thereafter, they were rinsed 3 times in 0.2% PBT. The samples were then blocked for 15 minutes in 5% Bovine Serum Albumin (BSA) (Roche, #10735094001) in 0.2% PBT. Next, the, samples were incubated in the primary antibody in 100 μl 5% BSA in 0.2% PBT overnight at 4 while rotating. The following primary antibodies were used: mouse anti-Abd-B (1/50, #1A2E9, DSHB), rabbit anti-P-AMPKalpha (T172) (1/100, #2535, Cell Signaling Technology), rabbit anti-cleaved Caspase-3 (1/200, #9661, Cell Signaling Technology), rabbit anti-cleaved Dcp-1 (1/200, #9578, Cell Signaling Technology), rabbit anti P-4E-BP1 (T37/46) (1/250, #2855, Cell Signaling Technology). After incubation in primary antibody, the carcasses were rinsed three times in 0.2% PBT for 10 minutes before being incubated for 2 hours at room temperature with the secondary antibody in 100 μl 5% BSA in 0.2% PBT. The following secondary antibodies used in this study: Alexa Fluor® 647 donkey-anti-rabbit (1:500, #715-605-152, Jackson), Alexa Fluo® 647 donkey anti-mouse (1:500, #715-605-150, Jackson). When necessary, phalloidin conjugated to Alexa Fluor™ 647 (Invitrogen^TM^, Cat; A22287, Lot: 1884190, dilution: 1:500 in 5% BSA in 0.2% PBT) was used for visualization of F-actin. Either Hoechst or DAPI was used to visualize the nuclei. If using Hoechst, the carcasses were stained with Hoechst 33342 (ThermoFisher, #62249) diluted to a concentration of 1 μg/ml in 0.2% PBT for 10 minutes. Then the carcasses were rinsed two times in 0.2% PBT for 10 minutes. The fat body was then separated from the rest of the carcass in a drop of PBS and mounted unto a microscope slide with ProLong™ Glass Antifade Mountant (Invitrogen™, #P36980). If Hoechst was not used to visualize the nuclei, ProLong™ Diamond Antifade with DAPI (Invitrogen™ #P36971) was used as the mounting media. The microscope slides were kept at 4 °C in the dark until the imaging session. Fluorescent imaging in absence of antibodies followed the same protocol, apart from blocking and staining with primary and secondary antibodies.

Imaging was performed using either the Andor Dragonfly 505 Spinning Disk confocal microscope (Oxford Instrumentals) equipped with an Andor Zyla 4.2 (sCMOS) camera and 405/ 488/561/637 nm lasers, using a Nikon CFI Plan Flour 40x (NA 1.30, immersion oil) objective and Fusion software. Alternatively, imaging was performed using the Nikon ECLIpS Ti-2E microscope equipped with a Yokogawa CSU-W1 spinning disk confocal unit, laser unit with 405/488/561/638 nm lasers, two sCOMS cameras (PrimeBSI) using the CFI Plan Apo λ 40x (NA 0.95, Air) objective and NIS-Elements 5.30 software. Alternatively, Nikon ECLIPSE Ti-2-E inverted microscope armed with CrestOptics X-Light V3 spinning-disk confocal module featuring 405/477/546/638 nm lasers was used. The samples were imaged using Plan Apo 60x NA 1.42 Oil λD objective. All confocal images in this study are maximum intensity projections of Z-stacks. Within each set of experiments, images were captured with identical settings below pixel value saturation and post-processed identically. Post-imaging enhancement of the fluorescent signal for adequate visualization was performed in ImageJ or NIS Elements (v 5.30).

### Image analysis

To measure GFP intensity of the fat body of whole larvae, larvae were segmented manually before measuring intensity of GFP for each larvae using ImageJ. Quantification of cell size, nuclei size and fluorescent signal intensities was performed in ImageJ or NIS elements. For mosaic images, the tissue, clones, and nuclei were found using an intensity-based approach on the mCherryAtg8a, GFP and DAPI channel, respectively. The mathematical approach Voronoi was employed to segment cells based on nuclei position relative to each other. Cells and nuclei touching image edges were discarded from the quantification. When appropriate, Atg8a spots were counted, and their size measured using an intensity-based approach on the mCherryAtg8a channel. For quantification of the double-tag, the presence of GFP in segmented mCherryAtg8a-spots was determined by appropriate thresholding on the GFP channel. PNGs with masks showing quantified cells and nuclei were saved to serve as a control for appropriate quantification. Data from ImageJ quantification was preprocessed in R/R Studio before visualization and statistical analysis were performed in GraphPad Prism 10.

### RT-qPCR

To extract RNA from the fat body of Drosophila, the larvae were rinsed in 0.2% Triton X-100 in 1X PBS (0.2% PBT) before dissection in ice cold 1X PBS. For each sample, fat bodies were collected in 80 µl 1X PBS then prepared for RNA extraction by grinding the samples for 30 seconds using a pestle motor. To isolate the RNA from the samples, TRIzol (Life Technologies, Cat#15596026, Lot#16200701) and the Direct-zol RNA Microprep kit (Zymo Research # R2062 # 236197) were employed, following the manufacturers provided protocol. The amount and quality of extracted RNA in nuclease free H2O was measured using the NanoDrop Spectrophotometer (Thermo Fisher). The iScriptTM cDNA Synthesis Kit (Bio-Rad #1708891 #644805223) was then incorporated for cDNA synthesis of the isolated RNA through reverse transcription with equal amounts of RNA from each sample. The cDNA was 5-fold diluted in nuclease free H2O before performing RT-qPCR. 2x Fast SYBR Green Mastermix (Applied Biosystems by Thermo Fisher Scientific. # 4385614) was mixed with 10 µl of primer solution containing the forward and reverse primers, nuclease free H_2_O and cDNA. The reaction was run using the QuantStudio™ 3 System (Applied Biosystems by Thermo Fisher Scientific) or Applied Biosystems StepOnePlus Real Time PCR system with StepOne v2.3 software. Rpl32 was employed as a housekeeping gene for normalization of gene expression levels within the same samples. qPCR primers were selected from http://www.flyrnai.org/flyprimerbank or custom designed. Primers used in this study are provided in Supplementary Table 2. Before using primers for the first time in the RT-qPCR assay, their efficiency and specificity of the were tested. To test the primer sets, cDNA was diluted in a 1:4 serial dilution of 4 dilutions to generate a four-point standard curve. The quality of the primer sets was estimated according to the melt curve and amplification efficiency.

### Statistical analysis

Samples were not masked or randomized during data collection or analysis. The statistical analysis in this study was performed in R or Graphpad Prism 10. Unless stated otherwise: The normality of the distribution in the dataset was tested by performing a Shapiro-Wilk test. If the data met the assumptions of normality, students t-test were performed to investigate differences between two groups. Ordinary one-way ANOVA with Dunnett’s post hoc test was performed to investigate differences between 3 or more groups. If assumptions of equal variances were not met despite normality in the data, Brown-Forsythe ANOVA was performed. If the data did not meet the assumptions of normality, the non-parametric Mann-Whitney U test was performed to investigate differences between two groups. Kruskal-Wallis test was performed to investigate significant differences between more than two independent groups followed by a Dunn’s post hoc test to identify which groups that differ from one another. The significant level of all tests was 0.05, meaning that p-values less than 0.05 was considered statistically significant (* p < 0.05, ** p < 0.01, *** p < 0.001, **** p < 0.0001).

## Acknowledgments

The authors would like to thank Renato Paro for generously sharing the MLL-AF4 fly lines. Stocks obtained from the Bloomington Drosophila Stock Center (NIH P40OD018537) and Vienna Drosophila Resource Center (VDRC) were used in this study. Several antibodies were obtained from the Developmental Studies Hybridoma Bank created by the NICHD of the NIH and maintained at The University of Iowa.

The MolMed Imaging Platform (MIP) at the Faculty of Medicine, Gaustad and the core facilities for Advanced Light Microscopy at Oslo University Hospital Montebello and Gaustad nodes, are acknowledged for access, help and services.

H.K. was supported by grants 2017062 and 2022019 from the South-Eastern Norway Regional Health Authorities, grant 300748 from the Research Council of Norway and funding from the European Union (ERC, FINALphagy, 101039174). The research leading to these results has received funding from the European Union’s Horizon 2020 research and innovation programme under the Marie Skłodowska-Curie grant agreement No 801133. This work was partly supported by the Research Council of Norway through its Centres of Excellence funding scheme, project number 262652. J.M.E. is supported by the Norwegian Health Authority South-East, grant numbers 2017064, 2018012 and 2019096; the Norwegian Cancer Society, grant numbers 182524 and 208012; and the Research Council of Norway grant numbers 261936 and 294916. Funded by the European Union. Views and opinions expressed are however those of the author(s) only and do not necessarily reflect those of the European Union or the European Research Council Executive Agency. Neither the European Union nor the granting authority can be held responsible for them.

## Author contributions

Aina Louise Cimafranca Haukeland: Formal analysis, Investigation, Writing—original draft, Writing—review & editing, Visualization.

Julie Aarmo Johannessen: Formal analysis, Investigation, Writing—review & editing, Visualization.

Nora Rojahn Bråthen: Formal analysis, Investigation, Writing—review & editing, Visualization.

Paul Bergeron: Formal analysis, Investigation, Writing—review & editing, Visualization.

Miriam Formica: Investigation, Writing—review & editing.

Jorrit M. Enserink: Writing—review & editing.

Helene Knævelsrud: Conceptualization, Formal analysis, Investigation, Writing—original draft, Writing—review & editing, Supervision, Project administration, Funding acquisition.

## Abbreviations

4E-BP1: Eukaryotic translation initiation factor 4E-binding protein
AMPK: AMP-activated protein kinase
Atg: Autophagy-related
CQ: Chloroquine
DIAP1: Drosophila inhibitor of apoptosis protein 1
ear: ENL/AF9-related
MLL: Mixed Lineage Leukemia
MLLr: MLL-rearranged
mTORC1: Mechanistic target of rapamycin complex 1
RNAi: RNA interference.

