## Supplemental figures for "The human leukemic fusion protein MLL-AF4 promotes autophagy and cell death in the fat body of *Drosophila melanogaster*"

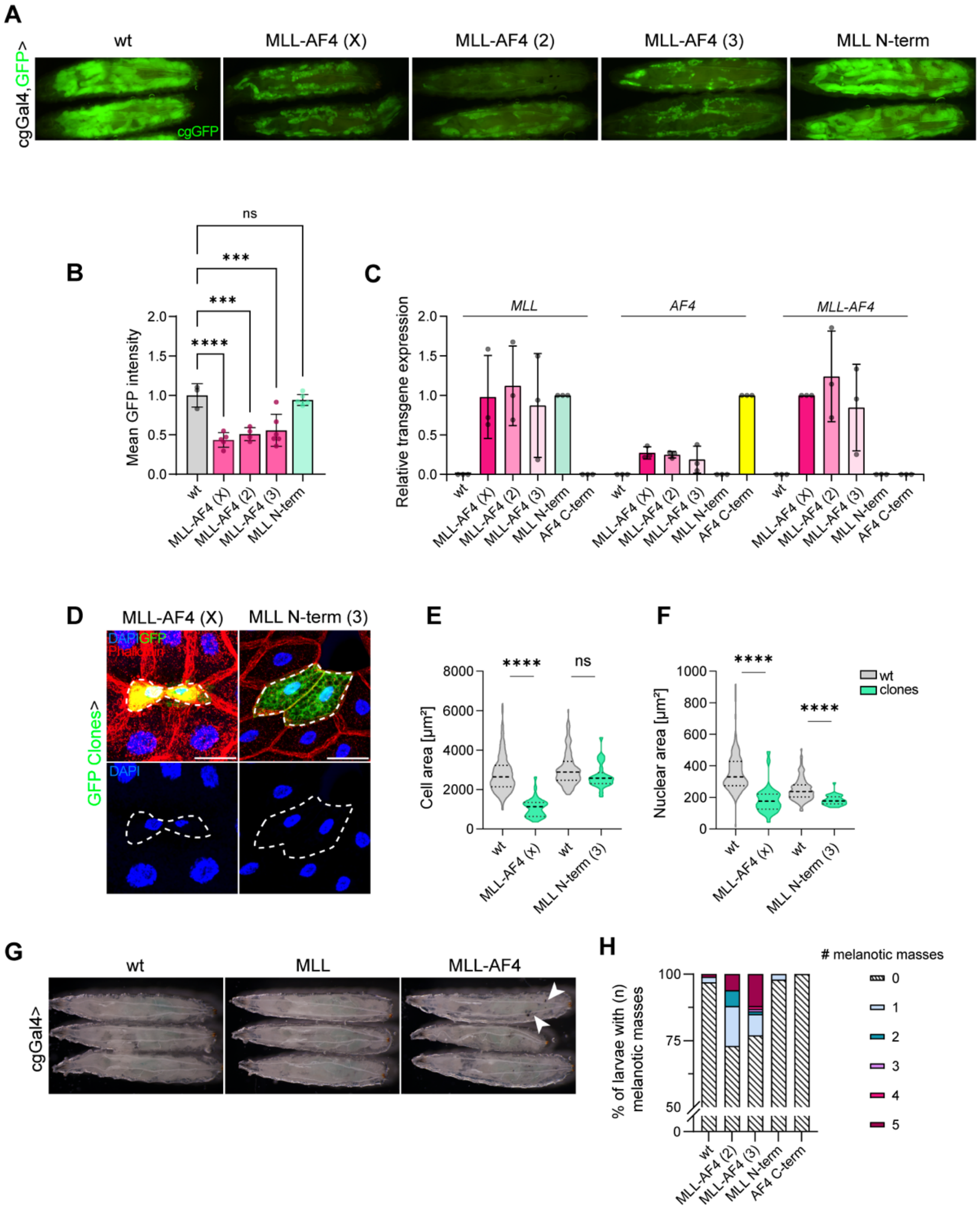


**Supplementary Figure 1. (A)** Wandering 3rd instar (wL3) larvae with cgGal4 driven expression of GFP and respective transgenes in the fat body. **(B)** Quantification of mean GFP intensity normalized to mean of wt. Graph shows mean ± SD. n = 3-6 larvae. **(C)** Relative gene expression levels of MLL, AF4 or MLL-AF4 as determined by qRT-PCR. Results are presented as the mean ± SD. **(D)** Mosaic fat body from wL3 larvae expressing GFP and MLL-AF4 or MLL N-terminus in clones. Scale bars = 50 µm. **(E)** Quantification of cell area of wt and clones from images in (D). **(F)** Quantification of nuclear area of wt and clones from images in (D). **(G)** wL3 larvae with respective genotypes expressed in the fat body. White arrows point to the presence of melanotic masses. **(H)** Bar plot shows percentage of larvae with (n) melanotic masses. MLL N-term, MLL N-terminus. wt, wild type.

Genotypes: (A, B) wt: *cgGal4, UAS-GFP/+*. MLL-AF4 (X): *UAS-MLL-AF4/+;cgGal4, UAS-GFP/+*. MLL-AF4 (2): *cgGal4, UAS-GFP/UAS-MLL-AF4.* MLL-AF4 (3): *cgGal4, UAS-GFP/+, UAS-MLL-AF4/+.* MLL N-term: *cgGal4, UAS-GFP/+, UAS-MLL N-terminal/+.* (C) wt: *cgGal4, UAS-GFP/+.* MLL-AF4 (X): *UAS-*

*MLL-AF4/+;cgGal4, UAS-GFP/+*. MLL-AF4 (2): *cgGal4, UAS-GFP/UAS-MLL-AF4.* MLL-AF4 (3): *cgGal4, UAS-GFP/+, UAS-MLL-AF4/+.* MLL N-term: *cgGal4, UAS-GFP/+, UAS-MLL N-terminal/+.* AF4 C-term: *cgGal4, UAS-GFP/+, UAS-AF4 C-terminal/+.* (D, E, F) MLL-AF4 (X): *hsflp/UAS-MLL-AF4;3xmCherryAtg8a, UAS-GFP/+;Act>>Gal4, UAS-Dcr2/+.* MLL-N-term (3): *hsflp/+;3xmCherryAtg8a, UAS-GFP/+;Act>>Gal4, UAS-Dcr2/UAS-MLL N-terminal.* (G, H) wt: *cgGal4, UAS-GFP/+.* MLL-AF4 (2): *cgGal4, UAS-GFP/UAS-MLL-AF4.* MLL-AF4 (3): *cgGal4, UAS-GFP/+, UAS-MLL-AF4/+.* MLL N-term: *cgGal4, UAS-GFP/+, UAS-MLL N-terminal/+.* AF4 C-term: *cgGal4, UAS-GFP/+,*

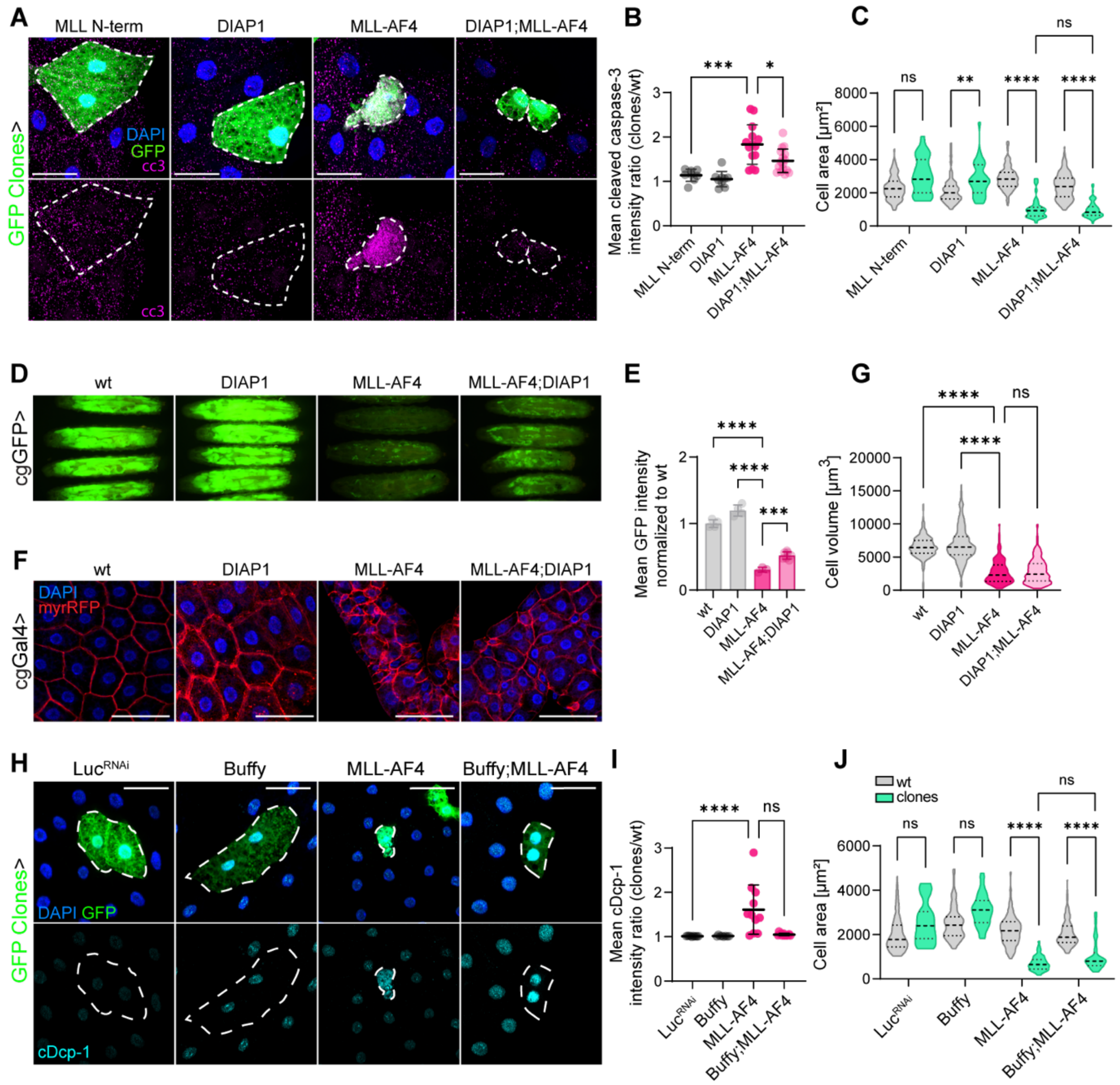


**Supplementary Figure 2.** **(A)** Mosaic fat body from wL3 larvae expressing GFP and respective transgenes in clones. Tissue is stained with antibody against cleaved caspase-3 to detect caspase activity. Scale bars = 50 µm. **(B)** Quantification of mean cleaved caspase-3 antibody intensity presented as the ratio between clones and wt. Graph shows mean ± SD. **(C)** Violin plot shows quantification of cell area of wt and clones. **(D)**. wL3 larvae with cgGal4 driven expression of GFP and respective transgenes in the fat body. **(E)** Quantification of mean GFP intensity normalized to wt. Graph shows mean ± SD. **(F)** Fat body from wL3 larvae expressing cgGal4 driven transgenes and myrRFP to visualize membranes. Scale bars = 100 µm. **(G)** Quantification of cell volume from images in **(F)**. **(H)** Mosaic fat body from wL3 larvae expressing GFP and respective transgenes in clones. Tissue is stained with antibody against cDcp-1 to detect caspase activity. Scale bars = 50 µm. **(I)** Quantification of mean cytoplasmic cDcp-1 intensity presented as the ratio between clones and wt. Graph shows mean ± SD. **(J)** Violin plot shows quantification of cell area of wt and clones from images in **(H)**. cc3, cleaved caspase-3. cDcp-1, cleaved Death caspase-1. MLL N-term, MLL N-terminus. wt, wild type.

Genotypes: (A, B, C) $Luc^{RNAi}$: *hsflp/+;3xmCherryAtg8a, UAS-GFP/+;Act>>Gal4, UAS-Dcr2/UAS-Dcr2/UAS-Luciferase-RNAi TRiP.JF01355*. DIAP1: *hsflp/+;3xmCherryAtg8a, UAS-GFP/ UAS-DIAP1.H (BL6657); Act>>Gal4, UAS-Dcr2/+.* MLL-AF4: *hsflp/+;3xmCherryAtg8a, UAS-GFP/ VDRCsh60200;Act>>Gal4, UAS-Dcr2/UAS-MLL-AF4.* DIAP1;MLL-AF4: *hsflp/+;3xmCherryAtg8a, UAS-GFP/UAS-DIAP1.H (BL6657); Act>>Gal4, UAS-Dcr2/ UAS-MLL-AF4.* (D, E) WT: *cgGal4, UAS-GFP/+.* DIAP1: *cgGal4, UAS-GFP/UAS-DIAP1 (BL6657).* MLL-AF4: *cgGal4, UAS-GFP/UAS-MLL-AF4.*

MLL-AF4;DIAP1 (3): *cgGal4, UAS-GFP/*+, *UAS-MLL-AF4/*+. MLL N-term: *cgGal4, UAS-GFP/UAS-MLL-AF4, UAS-DIAP1 (BL6657)/+.* (F, G) WT: *y,w, hsflp/+; cg-Gal4,FRT42D,UAS-myrRFP/+;UAS-GFP-Atg8a/*+, DIAP1: *y,w, hsflp/+; cg-Gal4,FRT42D,UAS-myrRFP/UAS-DIAP1 (BL6657);UAS-GFP-Atg8a/*+, MLL-AF4: *y,w, hsflp/+; cg-Gal4,FRT42D,UAS-myrRFP/UAS-MLL-AF4;UAS-GFP-Atg8a/*+, MLL-AF4;DIAP1: *y,w, hsflp/+; cg-Gal4,FRT42D,UAS-myrRFP/+;UAS-GFP-Atg8a/UAS-MLL-AF4; UAS-DIAP1 (BL6657)/*+. (H, I, J) Luc$^{RNAi}$: *hsflp/+;3xmCherryAtg8a, UAS-GFP/+;Act>>Gal4, UAS-Dcr2/ UAS-Luciferase-RNAi TRiP.JF01355*, Buffy: *hsflp/+;3xmCherryAtg8a, UAS-GFP/ UAS-Buffy (BL58358);Act>>Gal4, UAS-Dcr2/*+. MLL-AF4: *hsflp/+;3xmCherryAtg8a, UAS-GFP/40D-UAS;Act>>Gal4, UAS-Dcr2/UAS-MLL-AF4.* Buffy;MLL-AF4: *hsflp/+;3xmCherryAtg8a, UAS-GFP/ UAS-Buffy (BL58358) ;Act>>Gal4, UAS-Dcr2/UAS-MLL-AF4.*

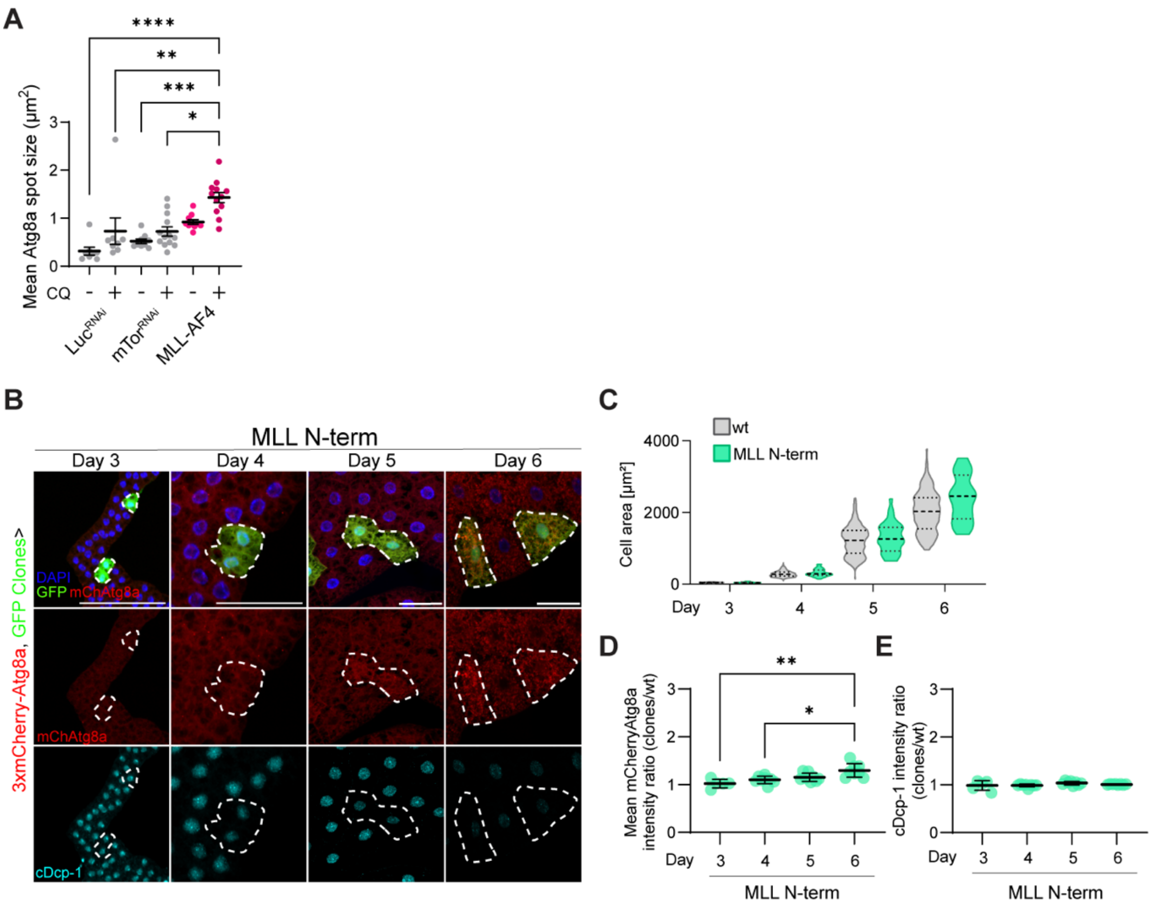


**Supplementary Figure 3. (A)** Quantification of mCherry-Atg8a spot size shown as the mean per cell per tissue section from images in Figure 3, C. **(B)** Mosaic fat body with GFP and MLL N-terminal moeity expressed in clones with endogenous 3xmCherry-Atg8a in the whole animal. Larvae were dissected in 24 hours intervals from day 3 to day 6 after eggs deposition. The tissue is stained with an antibody against cDcp-1 for caspase activity detection. Scale bars = 50 µm. **(C)** Quantification of cell area from images in (B). **(D)** Quantification of mean mCherryAtg8a intensity shown as a ratio between clones and wt cells. **(E)** Quantification of mean cytosolic cDcp-1 intensity shown as a ratio between clones and wt cells. (A, D, E) n = tissue sections. For appropriate panels, results are presented as the mean ± SD. cDcp-1, cleaved Death caspase-1. MLL N-term, MLL N-terminus. wt, wild type.

Genotypes: (A) Luc$^{RNAi}$: *cg-Gal4, UASp-GFP-mCherry-Atg8a/+;UAS-RNAi Luciferase TRiP.JF01355/+.* mTor$^{RNAi}$: *cg-Gal4, UASp-GFP-mCherry-Atg8a/+;UAS-RNAi mTor TRiP.HMS01114/+.* MLL-AF4: *cg-Gal4, UASp-GFP-mCherry-Atg8a/+;UAS-MLL-AF4/+.* (B, C, D, E) *hsflp/+;3xmCherry-Atg8a, UAS-GFP/+;Act>>Gal4, UAS-Dcr2/UAS-MLL N-terminal.*

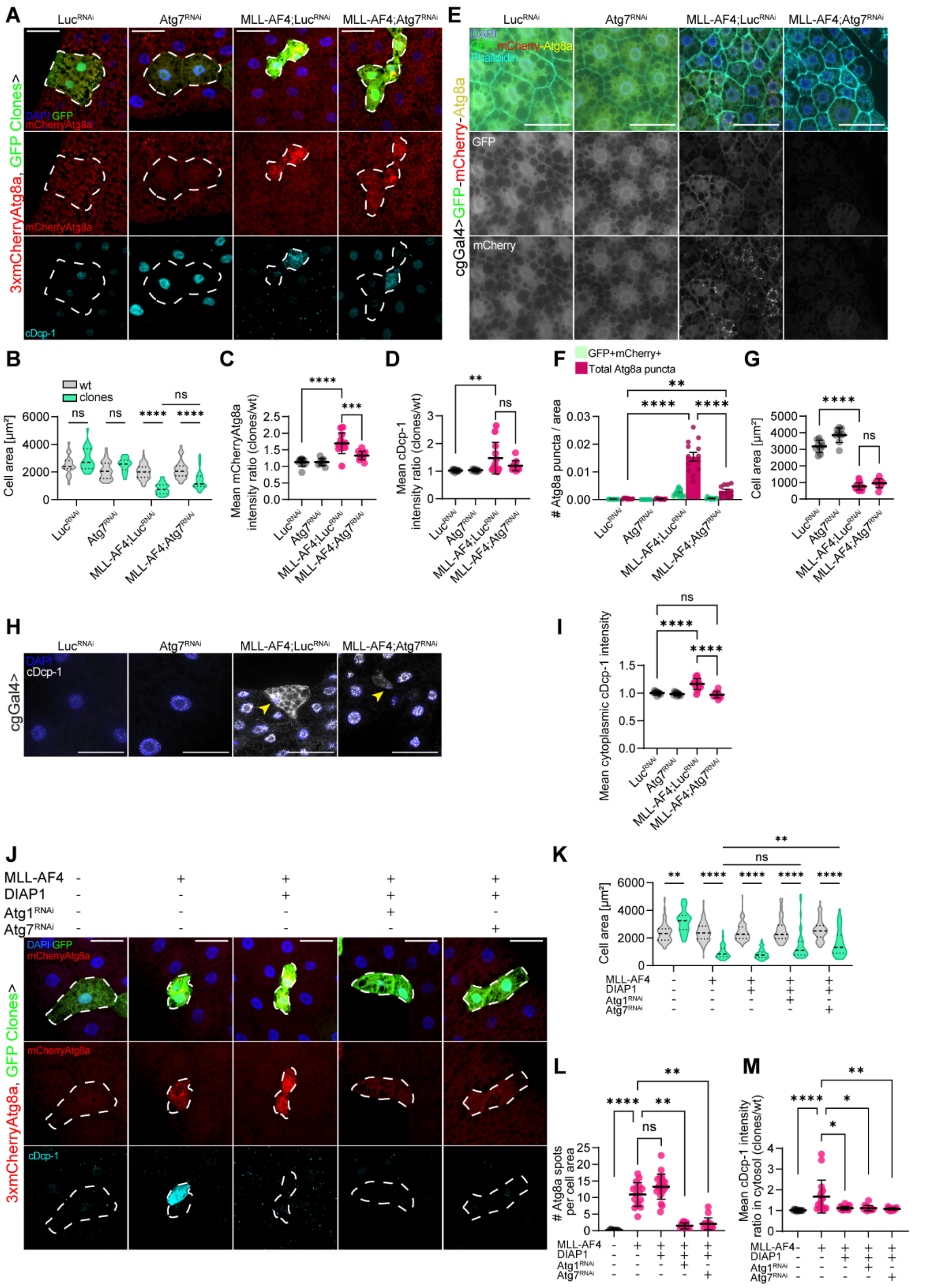


**Supplementary Figure 4. (A)** Representative confocal immunofluorescent images of wL3 mosaic fat body expressing respective transgenes in clones. Clones are outlined with a dotted line and express Gal4-driven GFP. The autophagy marker 3xmCherry-tagged Atg8a is expressed at endogenous levels in the whole animal. The tissue is stained with an antibody against cDcp-1. Scale bars = 50 µm. **(B)**

Quantification of cell area of images in (A). **(C-D)** Quantification of mean mCherryAtg8a (C) or cDcp-1 (D) intensity ratio between wt and clones from images in (A). **(E)** Representative confocal fluorescent images of fL3 fat body expressing respective transgenes and GFP-mCherry-Atg8a in the whole tissue. Scale bars = 50 µm. **(F)** Quantification of Atg8a puncta. Green bars represent Atg8a puncta positive for both GFP and mCherry, red bars represent total Atg8a puncta. **(G)** Quantification of mean cell area per tissue section. **(H)** Representative confocal immunofluorescent images of wL3 fat body expressing respective transgenes. The tissue is stained with an antibody against cDcp-1. Yellow arrowheads annotate cells with small nuclei and high cDcp-1 intensity. **(I)** Quantification of mean cDcp-1 intensity in cytoplasm normalized to mean of control. **(J)** Representative confocal immunofluorescent images of wL3 mosaic fat body expressing respective transgenes in clones. Clones are outlined with a dotted line and express Gal4-driven GFP. The autophagy marker 3xmCherry-tagged Atg8a is expressed at endogenous levels in the whole animal. The tissue is stained with an antibody against cDcp-1. **(K)** Quantification of cell area of images in (E). **(L)** Mean number of mCherryAtg8a positive puncta per cell normalized to cell area. (E). (**M**) Quantification of cDcp-1 intensity ratio between wt and clones from images in (J). For appropriate panels, results are presented as the mean ± SD. cDcp-1, cleaved Death caspase-1. fL3, feeding 3$^{rd}$ instar. wL3, wandering 3$^{rd}$ instar.

Genotypes: (A, B, C, D) Luc$^{RNAi}$: *hsflp/+;3xmCherryAtg8a, UAS-GFP/+;Act>>Gal4, UAS-Dcr2/UAS-Luciferase-RNAi TRiP.JF01355.* Atg7$^{RNAi}$: *hsflp/+;3xmCherryAtg8a, UAS-GFP/+;Act>>Gal4, UAS-Dcr2/ UAS-Atg7-RNAi TRiP.JF02787.* MLL-AF4;Luc$^{RNAi}$: *hsflp/+;3xmCherryAtg8a, UAS-GFP/UAS-MLL-AF4;Act>>Gal4, UAS-Dcr2/UAS-Luciferase-RNAi TRiP.JF01355.* MLL-AF4;Atg7$^{RNAi}$: *hsflp/+;3xmCherryAtg8a, UAS-GFP/UAS-MLL-AF4;Act>>Gal4, UAS-Dcr2/ UAS-Atg7-RNAi TRiP.JF02787.* (E, F, G, H I) Luc$^{RNAi}$: *cg-Gal4, UASp-GFP-mCherry-Atg8a/+;UAS-RNAi Luciferase TRiP.JF01355/+.* Atg7$^{RNAi}$: *cg-Gal4, UASp-GFP-mCherry-Atg8a/+; UAS-Atg7-RNAi TRiP.JF02787/+.* MLL-AF4;Luc$^{RNAi}$: *cg-Gal4, UASp-GFP-mCherry-Atg8a/UAS-MLL-AF4;UAS-RNAi Luciferase TRiP.JF01355/+.* MLL-AF4;Atg7$^{RNAi}$: *cg-Gal4, UASp-GFP-mCherry-Atg8a/UAS-MLL-AF4; UAS-Atg7-RNAi TRiP.JF02787/+.* (J, K, L, M) Control: *hsflp/+;3xmCherryAtg8a, UAS-GFP/+;Act>>Gal4, UAS-Dcr2/UAS-Luciferase-RNAi TRiP.JF01355.* MLL-AF4;Luc$^{RNAi}$: *hsflp/+;3xmCherryAtg8a, UAS-GFP/UAS-MLL-AF4;Act>>Gal4, UAS-Dcr2/ UAS-Luciferase-RNAi TRiP.JF01355.* MLL-AF4,DIAP1: *hsflp/+;3xmCherryAtg8a, UAS-GFP/UAS-MLL-AF4, UAS-DIAP1 (BL6657);Act>>Gal4, UAS-Dcr2/+.* MLL-AF4, DIAP1; Atg1$^{RNAi}$: *hsflp/+;3xmCherryAtg8a, UAS-GFP/UAS-MLL-AF4, UAS-DIAP1 (BL6657);Act>>Gal4, UAS-Dcr2/ UAS-Atg1-RNAi TRiP.JF02273.* MLL-AF4, DIAP1; Atg7$^{RNAi}$: *hsflp/+;3xmCherryAtg8a, UAS-GFP/UAS-MLL-AF4, UAS-DIAP1 (BL6657);Act>>Gal4, UAS-Dcr2/ UAS-Atg7-RNAi TRiP.JF02787*

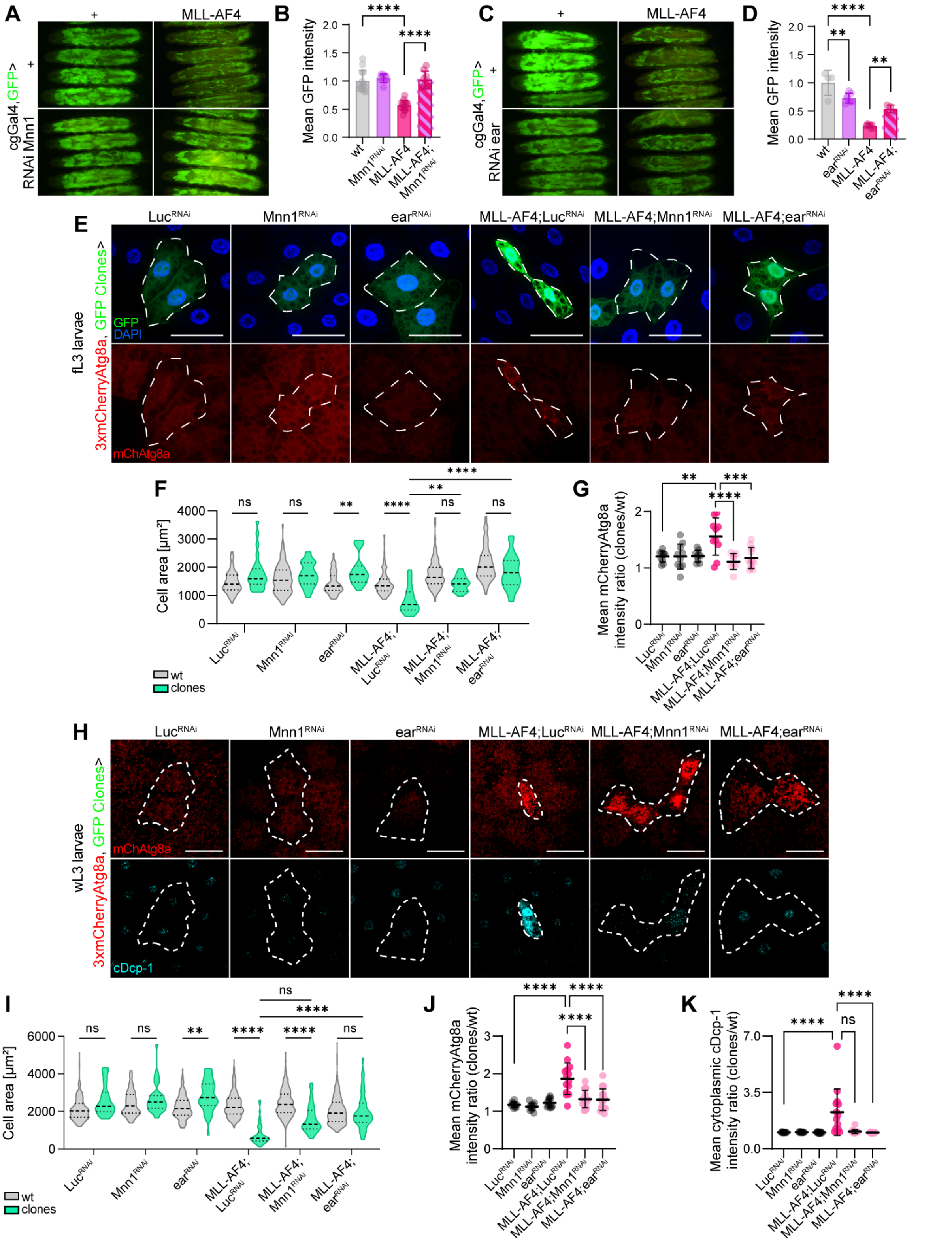


**Supplementary Figure 5. (A)** wL3 larvae with cgGal4 driven expression of GFP and respective transgenes in the fat body. **(B)** Quantification of mean GFP intensity normalized to wt. Graph shows mean ± SD, n = larvae. **(C)** wL3 larvae with cgGal4 driven expression of GFP and respective transgenes in the fat body. **(D)** Quantification of mean GFP intensity normalized to wt. Graph shows mean ± SD, n = larvae. **(E)** Mosaic fat body from fL3 larvae with GFP and respective transgenes expressed in clones and endogenous 3xmCherryAtg8a in the whole animal. **(F)** Quantification of cell area from images in (E). **(G)** Quantification of mCherryAtg8a intensity shown as a ratio between clones and wt cells. **(H)**

Mosaic fat body from wL3 larvae with GFP and respective transgenes expressed in clones and endogenous 3xmCherryAtg8a in the whole animal. The tissue is stained with an antibody against cDcp-1 for caspase activity detection. **(I)** Quantification of cell area from images in (H). **(J-K)** Quantification of mean mCherryAtg8a intensity (J) or cDcp-1 intensity (K) shown as a ratio between clones and wt cells. Scatterplot shows mean+-SD. Scale bars = 50 µm. cDcp-1, cleaved Death caspase-1. fL3, feeding 3$^{rd}$ instar. wL3, wandering 3$^{rd}$ instar. wt, wild type.

Genotypes: (A, B) wt: *cgGal4, UAS-GFP/+.* Mnn1: *cgGal4, UAS-GFP/+; RNAi Mnn1 TRiP.GL00018/+.* MLL-AF4: *cgGal4, UAS-GFP/UAS-MLL-AF4.* MLL-AF4;Mnn1 (3): *cgGal4, UAS-GFP/+, UAS-MLL-AF4/+.* MLL N-term: *cgGal4, UAS-GFP/UAS-MLL-AF4; RNAi Mnn1 TRiP.GL00018/+.* (C, D) WT: *cgGal4, UAS-GFP/+.* ear: *cgGal4, UAS-GFP/+; UAS-RNAi ear TRiP.JF02905/+.* MLL-AF4: *cgGal4, UAS-GFP/UAS-MLL-AF4.* MLL-AF4;ear (3): *cgGal4, UAS-GFP/+, UAS-MLL-AF4/+.* MLL N-term: *cgGal4, UAS-GFP/UAS-MLL-AF4; UAS-RNAi ear TRiP.JF02905/+.* (E-H) Luc$^{RNAi}$: *hsflp/+;3xmCherryAtg8a, UAS-GFP/+;Act>>Gal4, UAS-Dcr2/UAS-Luciferase-RNAi TRiP.JF01355.* Mnn1$^{RNAi}$: *hsflp/+;3xmCherryAtg8a, UAS-GFP/+;Act>>Gal4, UAS-Dcr2/ RNAi Mnn1 TRiP.GL00018/+.* ear$^{RNAi}$: *hsflp/+;3xmCherryAtg8a, UAS-GFP/+;Act>>Gal4, UAS-Dcr2/ UAS-RNAi ear TRiP.JF02905/+.* MLL-AF4;Luc$^{RNAi}$: *hsflp/+;3xmCherryAtg8a, UAS-GFP/UAS-MLL-AF4;Act>>Gal4, UAS-Dcr2/UAS-Luciferase-RNAi TRiP.JF01355.* MLL-AF4;Mnn1$^{RNAi}$: *hsflp/+;3xmCherryAtg8a, UAS-GFP/UAS-MLL-AF4;Act>>Gal4, UAS-Dcr2/ RNAi Mnn1 TRiP.GL00018.* MLL-AF4;ear$^{RNAi}$: *hsflp/+;3xmCherryAtg8a, UAS-GFP/UAS-MLL-AF4;Act>>Gal4, UAS-Dcr2/ UAS-RNAi ear TRiP.JF02905.*
