## Supplemental tables for "The human leukemic fusion protein MLL-AF4 promotes autophagy and cell death in the fat body of *Drosophila melanogaster*"

**Table S1:** Key resource table

| **REAGENT or RESOURCE** | SOURCE | IDENTIFIER |
| --- | --- | --- |
| **Primary Antibodies** | | |
| Mouse anti-Abd-B | DSHB | Cat# 1A2E9 |
| Rabbit anti-PAMPKalpha (T172) | Cell Signaling Technologies | Cat# 2535 |
| Rabbit anti Cleaved Caspase-3 (Asp175) | Cell Signaling Technologies | Cat# 9661 |
| Rabbit anti Cleaved Drosophila Dcp-1 (Asp215) | Cell Signaling Technologies | Cat# 9578 |
| Rabbit anti Phospho-4E-BP1 (Thr37/46) | Cell Signaling Technologies | Cat# 2855 |
| **Secondary Antibodies** |  |  |
| Alexa Fluor® 647 Donkey Anti-Rabbit | Jackson | Cat# 715-605-152 |
| Alexa Fluor^TM^ 647 Donkey Anti-Rabbit | Invitrogen Antibodies | Cat# A-31573 |
| Alexa Fluor® 647 Donkey Anti-Mouse | Jackson | Cat# 715-605-150 |
| **Chemicals, Peptides, and Recombinant Proteins** | | |
| Alexa Fluor^TM^ Phalloidin | Invitrogen™ | Cat# A22287 |
| Chloroquine diphosphate salt | Sigma-Aldrich | Cat# C6628 |
| Hoechst 33342 | ThermoFisher | Cat# #62249 |
| ProLong™ Glass Antifade Mountant | Invitrogen™ | Cat# P36980 |
| ProLong™ Diamond Antifade with DAPI | Invitrogen™ | Cat# P36971 |
| **Experimental Models: Organisms/Strains** | | |
| w, cg-Gal4, UAS-GFP |  |  |
| w[1118]; P{w[+mC]=Cg-GAL4.A}2 | BDSC | RRID:BDSC 7011 |
| y[1] w[1118]; P{w[+mC]=UASp-GFP-mCherry-Atg8a}2 | BDSC | 37749 |
| hsflp; 3xmCherry-Atg8a, UAS-GFP/CyO; Act>CD2>Gal4, UAS-Dicer2/TM6 | G Juhasz |  |
| w^iso^ | G. Laberge | N/A |
| w; UAS-MLL-AF4 ML5 (on 2^nd^) | R. Paro | N/A |
| w; UAS-MLL-AF4 ML6 (on 3^rd^) | R. Paro | N/A |
| w; UAS-MLL FL on 2^nd^ | R. Paro | N/A |
| w; UAS-MLL N-term (on 3^rd^) | R. Paro | N/A |
| w; UAS-MLL N-term (on X) | R. Paro | N/A |
| w; UAS-AF4 C-term on X | R. Paro | N/A |
| 40D-UAS | VDRC | 128207 |
| w[1]; P{w[+mC]=UAS-Abd-B.m.C}1.1 | BDSC | 913 |
| UAS-RNAi Abd-B TRiP.JF02309 | BDSC | 26746 |
| UAS-RNAi AMPKalpha TRiP.JF01951}attP2 | BDSC | RRID:BDSC 25931 |
| UAS-Atg1-RNAi TRiP.JF02273 | BDSC | RRID:BDSC 26731 |
| UAS-Atg7-RNAi TRiP.JF02787 | BDSC | RRID:BDSC 27707 |
| w[*]; P{w[+mC]=UAS-Buffy.Q}2 | BDSC | RRID:BDSC 58358 |
| w[*]; P{w[+mC]=UAS-DIAP1.H}3 | BDSC | RRID:BDSC 6657 |
| UAS-RNAi ear TRiP.JF02905}attP2 | BDSC | RRID:BDSC_28068 |
| y[1] v[1]; UAS-luciferase-RNAi TRiP.JF01355 | BDSC | RRID:BDSC_31603 |
| UAS-RNAi Mnn1 TRiP.GL00018}attP2 | BDSC | RRID:BDSC 35150 |
| UAS-RNAi Tor TRiP.HMS01114}attP2 | BDSC | RRID:BDSC 34639 |
| w[1118];P{VDRCsh60200}attP40 | VDRC | 60200 |
| **Oligonucleotides** | | |
| RT-qPCR primer sets | See Table S2 | N/A |
| **Software and Algorithms** | | |
| GraphPad Prism 10 | GraphPad Software | RRID:SCR_002798 |
| NIS Elements | Nikon | RRID:SCR_014329 |
| FIJI |  | RRID:SCR_002285 |
| Fusion Release 2.3 | Oxford Instruments |  |
| QuantStudioTM Design & Analysis Software v1.5.1 |  |  |
| StepOne software | Applied Biosystems | RRID:SCR_014281 |

**Table S2:** RT-qPCR primer sets

| **Target** | **Forward primer sequence** | **Reverse primer sequence** |
| --- | --- | --- |
| abd-A | ATCCCTGGATGACGCTTACAG | CGAGTGTAGGTCTGGCGAC |
| Abd-B | TGTTCGCCTATCCTTACCCAG | CAGAGGTGGTCTGATCGGG |
| AF4 C-term | CAGACTCCCATTGCCTTTGA | AGCAGGTCTAGGGTGATCTT |
| Atg8a | GGTCAGTTCTACTTCCTCATTCG | GATGTTCCTGGTACAGGGAGC |
| Dfd | CATCATGTCCAGAGTCCCATGT | GCGGTCATCGAATAGTGTCCATT |
| lab | AATGAAACGCAGGTCAAAATCTG | CTGTCAGTGTCATGGCTGTC |
| MLL-AF4 | ACTCCTAGTGAGCCCAAGAA | CTTATTGACCGGAGGTGGTTT |
| MLL | CTCCTCTCTTCCCTTGGTTTAC | CTCTTGTCAGCATCTCGATCTT |
| pb | GAGTCCCAAAATCGGAACACC | CGACGCCGACATTGACATTAAC |
| Rpl32 | GCCCAAGGGTATCGACAACA | GCGCTTGTTCGATCCGTAAC |
| Thor | CAGATGCCCGAGGTGTACTC | TTCATGAAAGCCCGCTCGTA |
| Ubx | GCATGAGTCCCTATGCCAACC | TCGCCGTATATCTTGTTGCAG |
